# Cell lineage as a predictor of immune response in neuroblastoma

**DOI:** 10.1101/2021.01.29.428154

**Authors:** Satyaki Sengupta, Sanjukta Das, Angela Crespo, Brian Miller, Bandana Sharma, Shupei Zhang, Ruben Dries, Hao Huang, Malgorzata Krajewska, David N. Debruyne, Luigi Soriano, Malkiel A. Cohen, Rogier Versteeg, Rudolf Jaenisch, Stefani Spranger, Judy Lieberman, Rani E. George

## Abstract

Immunotherapy for patients with neuroblastoma has met with limited success, partly due to an incomplete understanding of the mechanisms underlying immune responsiveness in this clinically and genetically heterogenic tumor. Here, we undertook an unbiased analysis using dimension reduction and UMAP visualization of transcriptional signatures derived from 498 primary neuroblastoma tumors. Four distinct clusters based on differentially expressed genes emerged, of which one, representing about 30% and comprising mainly of *MYCN*-nonamplified tumors, was notable for the high expression of genes associated with both immune response activation and suppression. This capacity to elicit a productive immune response resided exclusively in tumors with dominant populations of undifferentiated, neural crest-like or mesenchymal cells; by contrast, tumors comprising primarily of committed, adrenergic neuron-like cells were less immunogenic. Mesenchymal neuroblastoma cells were enriched for innate and adaptive immune gene signatures, demonstrated engagement with cytotoxic T and natural killer cells, and induced immune cell infiltration in an immunocompetent mouse model. Transcriptional or targeted therapy-induced reprogramming of adrenergic cells to the mesenchymal state led to reactivation of tumor cell-intrinsic immune genes. Key immune response genes in adrenergic tumor cells were found to be epigenetically silenced by the PRC2 complex, and such repression could be relieved by either mesenchymal cell state reprogramming or EZH2 inhibition, leading to increased activation of natural killer cells by the tumor cells. These data identify cell lineage as a major determinant of the immunogenic potential in neuroblastoma that could be used to stratify patients who are most likely to benefit from immunotherapy.

## INTRODUCTION

The anti-disialoganglioside GD2 monoclonal antibody dinutuximab has significantly improved event free survival rates in neuroblastoma^1^. Derived from the developing neural crest, this common solid tumor of childhood manifests as an extracranial mass arising in the adrenal medulla or sympathetic ganglia. Approximately half of all patients have high-risk features associated with a poor outcome - age >18 months, distant metastases and unfavorable histologic and genetic factors including amplification of the *MYCN* oncogene^2^. The success of anti-GD2 therapy that relies on immune cell-mediated cytotoxicity suggests that patients with neuroblastoma would benefit from other forms of immunotherapy; however, treatment results with use of cytotoxic CD8+ T lymphocytes directed against neuroblastoma antigens^3^, adoptive transfer of chimeric antigen receptor (CAR)-modified T cells^4–6^ or checkpoint inhibition^7,8^ have been suboptimal.

Major impediments to the effectiveness of immunotherapy in neuroblastoma are the immune evasion tactics deployed by the tumor cells as well as the tumor microenvironment (TME)^9^. These include downregulation of major histocompatibility complex (MHC) class I molecules and defects in antigen-processing machinery (APM) that render neuroblastoma cells resistant to T-cell-mediated cytotoxicity^10–12^, downregulation of cell-surface ligands required for natural killer (NK) cell receptor activation^13^, upregulation of checkpoint proteins that exert a protective role from NK cell-mediated lysis^14^, inefficient homing of cytotoxic T-cells to the tumor site^15^ or tumor cell overexpression of the leukocyte surface antigen CD47, which enables avoidance of macrophage-mediated phagocytosis^16^. Moreover, infiltration of suppressive immune cells such as T regulatory cells^17^, tumor-associated macrophages (TAMs)^18,19^, myeloid-derived suppressor cells^20^ and secreted immunosuppressive factors such as TGF-β, contribute to the generation of a TME that hinders an effective immune response and further dampens the effects of adoptive cell therapies^21^.

Amplification of the MYCN oncogene poses another distinct challenge to immunotherapy in neuroblastoma. This transcription factor is amplified in approximately 50% of high-risk cases and is associated with aggressive disease and a poor clinical outcome^22,23^. *MYCN*-amplified tumors consistently evade immune destruction by downregulating MHC class I molecules^10^ and are associated with poor infiltration of cytotoxic CD8^+^ T cells^24,25^ and reduced expression of NK cell ligands^26^. Interestingly, approximately half of high-risk neuroblastomas do not express amplified *MYCN*, and their capacity to induce a productive immune response remains unclear. In a recent study that analyzed the immune gene expression programs associated with *MYCN-*nonamplified tumors from high-risk patients, tumors with low as well as high functional tumor *MYCN* signatures were observed to have significantly higher levels of NK and CD8+ T-cell infiltrates compared to *MYCN*-amplified tumors; although, somewhat counterintuitively, these findings translated into a better outcome only in patients with high *MYCN* tumor signatures^25,27^.

Thus, although many of the mechanisms of immune evasion in neuroblastoma are known, further understanding of the tumor-host interaction will be crucial to enhancing the ability of immunotherapy to target and eliminate tumor-initiating and propagating cell populations. Especially challenging is the genetic and biologic heterogeneity of this tumor which makes it difficult to identify factors that consistently indicate the likelihood of an effective immune response and hence identify patients who are most likely to benefit from this form of therapy. Thus, we undertook an unbiased analysis of gene expression signatures across diverse clinical subtypes of primary tumors and identify tumor cell state as an important predictor of immune responsiveness in neuroblastoma.

## RESULTS

### A subset of primary neuroblastomas express markers of a productive immune response

To determine whether neuroblastomas are capable of eliciting a productive immune response, we first examined bulk RNA-sequencing data from 498 well-annotated primary human tumors representing diverse clinical and genetic subtypes (SEQC-498; GSE49711; **Supplementary Fig. 1a**) to quantify tumor-to-tumor gene expression variability and cluster tumor types based on gene expression profiles (see also Methods). In this unbiased analysis, all tumors within one cluster would share similar gene expression profiles, while being dissimilar to those of tumors within other clusters. Specifically, we first identified the top 5000 highly variably expressed genes within this dataset based on the premise that these would be most likely to contribute to distinct molecular subtypes^28,29^ (**Supplementary Fig. 1b; Supplementary Table 1)**. The data were dimensionally reduced using principal component analysis (PCA) and the top 20 leading principal components selected for clustering analysis (**Supplementary Fig. 1c**). Four distinct clusters were identified and visualized using 2D-Uniform Manifold Approximation and Projection (UMAP), a non-linear dimension-reduction tool^30,31^ (**Fig. 1a**). To explore the transcriptional differences between the clusters, we identified the differentially expressed genes (DEGs) in each cluster and noted that tumors in cluster 1 (C1; n = 103), termed *Hi-MYCN*, were enriched for MYCN target genes involved in cell proliferation and biosynthesis, and comprised 20% of the tumor set (**Fig. 1a-c; Supplementary Fig. 1d; Supplementary Table 2**). Not surprisingly, this cluster segregated with *MYCN-*amplified tumors in patients aged ≥ 18 months with stage 4 disease [according to the international neuroblastoma staging system (INSS)]^32^ and annotated “high risk” status (based on the Children’s Oncology Group risk classification)^33^ (**Fig. 1d; Supplementary Fig. 1e**). The remaining clusters consisted of *MYCN-*nonamplified tumors (**Fig. 1a**) of which, cluster 2 (C2, n = 241), or *neuronal*, made up the largest proportion of tumors, 48%, and comprised tumors that were enriched for DEGs with roles in nervous system development **Fig. 1a-c; Supplementary Fig. 1d**). Cluster 3 (C3, n = 140), accounting for 28% of the tumors, was enriched for tumors whose DEGs were involved in immune function, such as interferon-gamma (IFN-γ) response and T cell inflammation and activation, and hence were designated *immunogenic* (**Fig. 1a-c; Supplementary Fig. 1d**). Cluster 4 (C4; n=14; 3%) was clearly distinct from the other three clusters, and largely consisted of the spontaneously regressing stage 4S tumors that were predominantly enriched for genes involved in fatty acid and cholesterol homeostasis and hence were termed *metabolic* (**Fig. 1a-c; Supplementary Fig. 1d**). The neuronal and metabolic tumors arose predominantly in children <18 months of age, were of stages 1-3 and 4S, while the tumors within the immunogenic cluster were associated with patient age ≥18 months and metastatic disease (n= 66; 47%) (**Fig. 1d; Supplementary Fig. 1e**). Thus, our DEG-based analysis of almost 500 tumors categorized neuroblastoma into four largely distinct groups that included a distinct subset, accounting for approximately one-third of the entire cohort, whose gene expression profiles were closely linked to immune responsiveness. To ensure that these results were not confined to one data set, we analyzed an independent data set of 394 tumors (GSE120572) using similar clustering methods. This cohort also segregated into *Hi-MYCN, neuronal* and *immunogenic* clusters, again denoting the presence of immune response gene expression in a subset of primary neuroblastomas, the majority of which lack MYCN amplification (**Supplementary Fig. 1f, g**).

**Fig. 1.**
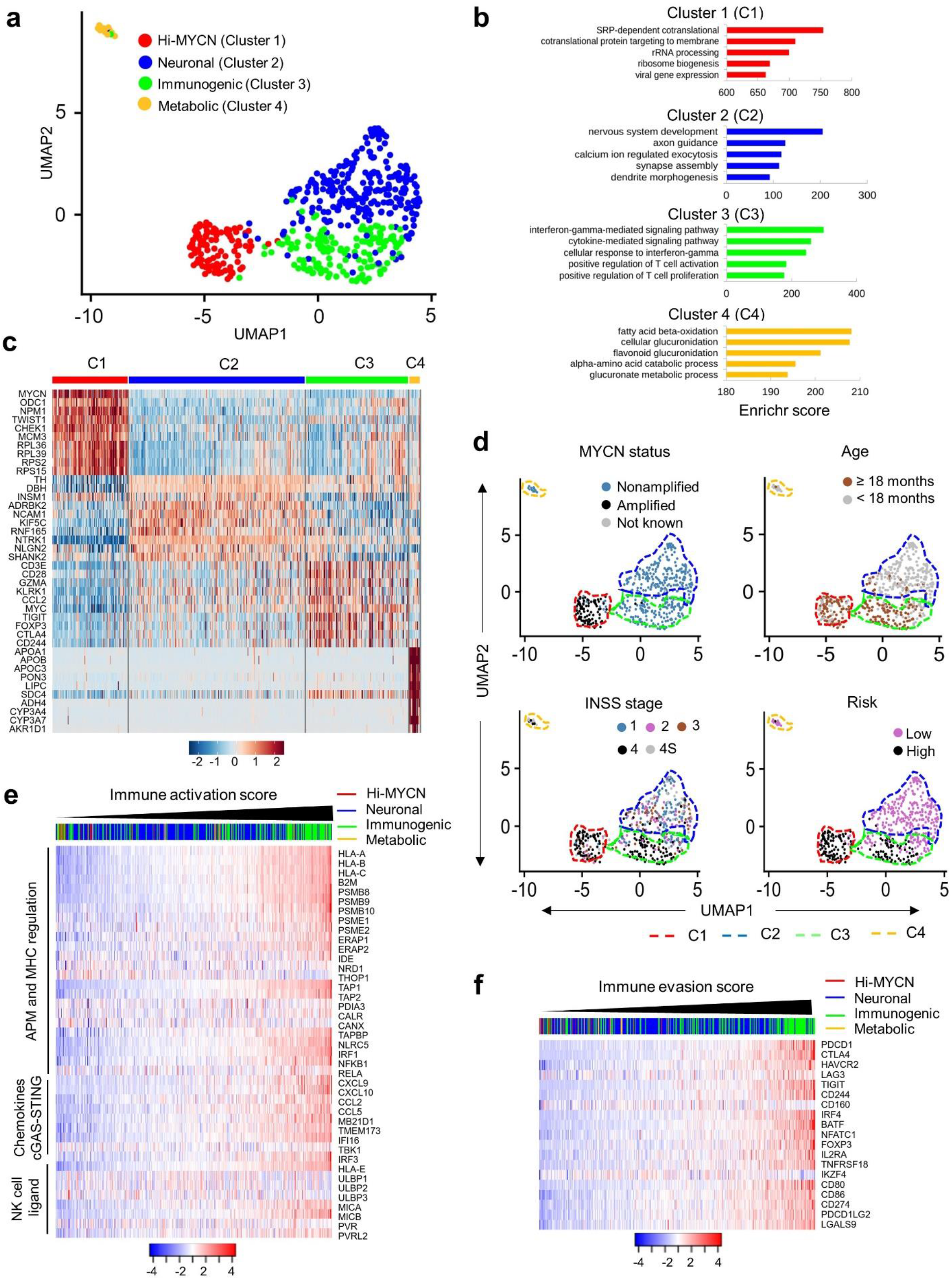
A subset of neuroblastomas exhibits a productive immune response. (**a**) Two-dimensional UMAP representations of the gene expression profiles in 498 neuroblastoma (NB) tumors. Each dot represents a tumor. The top 5000 highly variable genes were selected based on the variance-stabilizing method^34^ and the 20 significant principal components (PCs) selected and processed in UMAP to generate four clusters representing four NB subtypes. The DEGs were identified for each cluster using the receiver operating characteristics (ROC) curve to compare one cluster with other three (log_2_ FC > 0.25). (**b**) Gene ontology (GO) analysis of top DEGs in the four clusters. (**c**) Heat map of expression values of 10 representative DEGs within each cluster. Rows are z-score scaled average expression levels for each gene in all four clusters. (**d**) UMAP visualization of the distribution of the indicated prognostic features in NB among the four different clusters. (**e, f**) Heat map of z-score transformed log_2_ normalized expression values of immune activation (**e**) and evasion (**f**) genes in *MYCN-*nonamplified NBs (n=401). Tumors were ranked based on increasing immune activation or evasion scores. Cluster annotations of the tumors are indicated on the top horizontal bar.

To pursue the immune genes that were differentially enriched in the immunogenic cluster, we generated an immune activation (IA) score based on the relative expression of a curated set of 41 genes known to have major roles in tumor cell-intrinsic immune functions, such as regulation of MHC expression, antigen processing and presentation, NK cell recognition and T and NK cell infiltration (**Supplementary Table 3**). After assigning an IA score to each of the *MYCN-* nonamplified tumors (n = 401) in the SEQC-498 data set and arranging them in ascending order (**Supplementary Table 4**), we observed that a significant number with the highest IA scores predominantly fell within the immunogenic and metabolic clusters (**Fig. 1e; Supplementary Fig. 2a**), while those with intermediate or lower scores were associated with the neuronal and Hi-MYCN clusters, respectively (**Fig. 1e; Supplementary Fig. 2a**). Because a cytotoxic immune response is generally accompanied by immune suppression or evasion^35,36^, we determined whether immune suppression was also represented in the *MYCN-*nonamplified tumors by ranking them in ascending order of an immune evasion (IE) score based on the relative expression of 19 genes, most of which were markers of T-cell dysfunction (**Supplementary Table 3**). Again, the immunogenic tumor cluster had significantly higher IE scores compared with the neuronal and metabolic clusters (**Fig. 1f; Supplementary Fig. 2b**). Moreover, we observed enrichment for IA and IE scores in the immunogenic cluster in the additional data set (GSE120572) (**Supplementary Fig. 2c**), thus strengthening our premise that these tumors maybe capable of eliciting an immune response.

Consistent with the known poor immunogenicity of *MYCN*-amplified tumors^24,25^, we also observed that tumors within the Hi-MYCN cluster had, on the whole, the lowest IA and IE scores (**Supplementary Fig. 2a, b**). Surprisingly, however, a small subset within this cluster had scores that were comparable to the highly immunogenic tumors within the immunogenic cluster [13 of 103 (12.6%) above the median for immunogenic tumors] (**Supplementary Fig. 2a, b**). Thus, while the majority of neuroblastomas do not possess an immune response gene signature, a subset has significantly increased expression of both immune activation and evasion markers, pointing to their ability to induce an anti-tumor immune response.

### The mesenchymal lineage is preferentially associated with immune response signatures in neuroblastoma

Having identified subsets of neuroblastomas with the potential for immunogenicity, we next sought a biomarker that might consolidate the complex interactions between the immune system and the tumor. To this end, we performed a modular gene co-expression analysis of the 140 transcriptomes within the immunogenic cluster in the SEQC-498 data set to identify biologically relevant pathways based on similar gene expression patterns. Using the CEMiTool (co-expression modules identification tool) package (Russo et al., 2018), we identified five gene co-expression modules (M1-M5) within the immunological cluster (**Supplementary Fig. 3a**). Among these modules, M1, with the highest number of co-expressed genes, contained gene sets enriched for epithelial to mesenchymal transition (EMT), inflammatory response, and interferon signaling, suggesting an association between EMT and the preponderant representation of immune marker genes within the immunogenic cluster (**Fig. 2a; Supplementary Fig. 3b**). Furthermore, integration of the co-expression data in module M1 with protein–protein interaction data from the STRING 11.0 database identified mesenchymal lineage and immune markers as top regulatory hubs (**Fig. 2b**), leading us to hypothesize that in neuroblastoma, tumor immunogenicity could be determined by cell state.

**Fig. 2.**
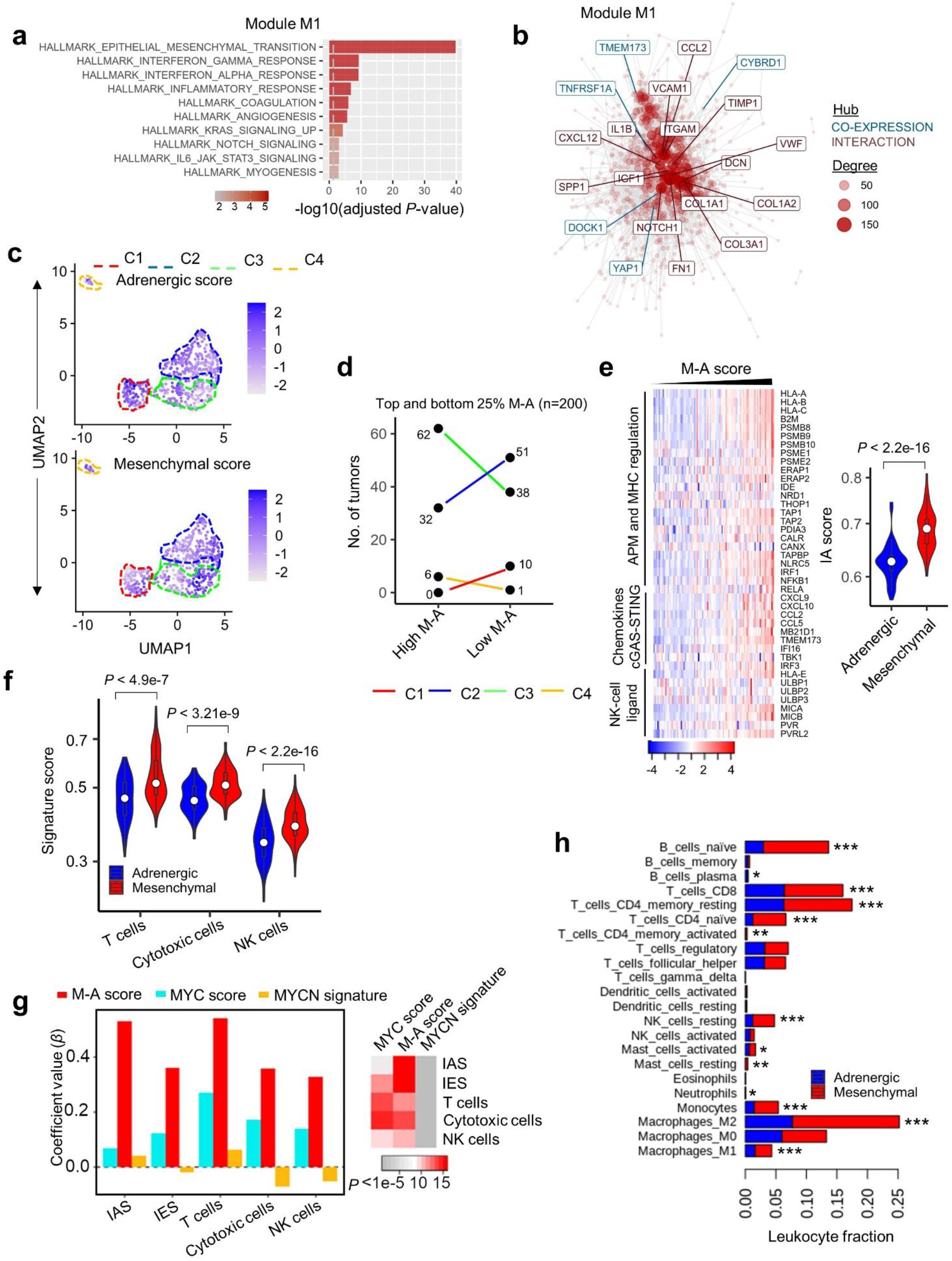
The mesenchymal cell state is associated with an immunogenic signature in NB. (**a**) GO analysis of co-expressed genes associated with module M1 using the KEGG (Kyoto encyclopedia of genes and genomes) database. The vertical dashed line indicates the adjusted *P*-value of 0.05. (**b**) Gene network representing all possible interactions in module M1. The topmost connected genes (hubs) are indicated. Hubs derived from module M1 are colored blue (co-expression) and those from the STRING database are indicated in red (interaction). The size of each node corresponds to the degree of interaction. (**c**) UMAP visualization of the distribution of adrenergic (top) and mesenchymal scores (bottom) among the four tumor clusters. Color bar represents normalized z-scores. Values <2.5 and >2.5 were set to −2.5 and +2.5 respectively, to reduce the effects of extreme outliers. (**d**) Dot plots showing the distribution of *MYCN-* nonamplified tumors (n = 400) within each of the clusters based on ranked M-A scores. *Left*, Tumors from the upper (high M-A) and lower (low M-A) M-A score quartiles are shown (n = 200; *P* < 0.01 for C3). *Right*, Representations based on the median M-A scores of the entire tumor cohort (n = 400; *P* = 0.05 for C3). Fisher’s exact test was used for both calculations. (**e**) *Left*, Heatmap representation of the expression of tumor cell-intrinsic immune activation genes in *MYCN-*amplified tumors (n=92). Samples are ranked by increasing M-A score. Log_2_ gene expression values were z-score transformed for heatmap visualization. *Right*, Violin plots of the distribution of immune activation scores in the tumors on the left, classified either as adrenergic or mesenchymal, based on the median M-A score. The box plots within the violin plots are defined by center lines (medians), box limits (25^th^ and 75^th^ percentiles), whiskers (minima and maxima; 1.5X the interquartile range). Significance was determined by the two-sided Kolmogorov-Smirnov (KS) test. APM, antigen processing machinery. (**f**) Violin plots comparing the quantitative scores of the indicated immune cell signatures in 100 tumors from the upper (mesenchymal) and lower (adrenergic) quartiles of the tumor M-A scores using the two-sided KS test. The box plots within the violin plots are defined as in (D). (**g**) *Left*, Bar diagram comparing regression coefficient (*β*) values derived from multivariate multiple regression model analysis of *MYCN-*nonamplified tumors. *β*-coefficient values were compared between three predictors: MYC score, M-A score and MYCN signature. IA, IE, T cell, cytotoxic cell and NK cell scores were used as response variables to generate the model^39^. *Right*, Heat map of the *P-*values associated with the three predictors. (**h**) Bar diagram comparing the CIBERSORT-estimated fractional content of the indicated tumor-infiltrating leukocytes between *MYCN-*nonamplified adrenergic and mesenchymal tumors. Adrenergic and mesenchymal tumors were assigned as in (**f**). Data represent the means, n = 100 tumors, *\*P* < 0.05, *\*\*P* < 0.01, *\*\*\*P* < 0.001 two-tailed Welch’s t-test.

Two independent groups ^37,38^ recently described two distinct cell states in neuroblastoma: a differentiated sympathetic neuron-like adrenergic (ADR) phenotype, defined by lineage markers including *PHOX2B, DBH*, and *TH*, and a mesenchymal (MES) phenotype, characterized as “neural crest cell-like” (NCC), and expressing genes such as *PRRX1, FOSL1*, and *FOSL2*. To test our prediction, we first quantified the adrenergic and mesenchymal identities of each tumor in our cohort based on the expression levels of the lineage-specific genes in each cell state as established by Groningen et al^38^ (see Methods). We ensured that there was no overlap between the 369-gene adrenergic signature and the genes that made up the IA data set and removed the 6 IA genes that were also present in the 485-gene mesenchymal signature. Next, we assigned either an adrenergic (A-score) or a mesenchymal (M-score) score to each tumor within our four previously identified clusters. This analysis revealed significant enrichment of the mesenchymal cell state within the immunogenic and metabolic clusters (**Fig. 2c; Supplementary Fig. 3c**). By contrast, the Hi-MYCN and neuronal clusters were enriched for the adrenergic cell state (**Fig. 2c; Supplementary Fig. 3c**).

Next, to identify predictors of immunogenicity among the *MYCN-*nonamplified tumors, we calculated the relative mesenchymal score for each tumor by subtracting the adrenergic from the mesenchymal score (M-A score). This resulted in a continuum of low to high M-A scores, corresponding to a less mesenchymal to a more mesenchymal tumor state (**Supplementary Fig. 3d**). To determine whether these cell states had any effect on immune response in the *MYCN-* nonamplified tumors, we determined whether the mesenchymal, adrenergic, or M-A scores correlated with our previously defined immune activation and evasion scores (**Fig. 1e, f**). We also tested the effect of *MYC*, one of the top differentially expressed genes in the immunogenic cluster (**Supplementary Fig. 3e**). *MYC* is overexpressed in approximately 10% of *MYCN-*nonamplified neuroblastomas^40^ and regulates the expression of cell-intrinsic immune evasion markers in lymphoma^41^. Of these variables, the M-A score showed the strongest correlation with immunogenicity, not only in terms of the immune activation score (R = 0.71), but also the immune evasion score (R = 0.51) (**Supplementary Fig. 3f**). MYC expression was only modestly correlated with immune evasion and activation scores (R = 0.43; R= 0.38, respectively) (**Supplementary Fig. 3g**). With no overlap between the lineage marker and immune response gene sets, these results suggest that the relative abundance of a mesenchymal signature (M-A score) is a better predictor of immune response than individual adrenergic or mesenchymal signatures. In agreement, tumors with high M-A scores were represented at a significantly higher proportion within the immunogenic cluster compared to tumors with low M-A scores (**Fig. 2d**), a result that was recapitulated in our second data set, (GSE120572) (**Supplementary Fig. 4a**). Finally, our finding of the subset of tumors marked by the relatively high expression of immune activation and evasion genes within the Hi-MYCN cluster (**Supplementary Fig. 2a, b**) prompted us to further evaluate the cell states of these tumors. Ranking these tumors based on increasing M-A scores revealed a positive relationship between cell-intrinsic immunogenicity and the mesenchymal state (**Fig. 2e**), suggesting that similar to our results in *MYCN-*nonamplified tumors, the presence of immune gene expression in *MYCN-*amplified tumors is significantly correlated with the mesenchymal phenotype.

The preferential overexpression in the mesenchymal phenotype of tumor cell-intrinsic genes that induce a positive immune response (IA score), as well as its correlation with transcripts that suppress the immune response (IE score), suggested that these tumors may support increased immune cell infiltration. To test this prediction, we used two orthogonal approaches. First, we assessed whether established signatures of immune cell infiltration ^39,42^ were present in the tumors arranged according to increasing M-A scores, and observed enrichment for signatures of infiltrating immune cells in tumors with mesenchymal phenotypes (**Supplementary Fig. 4b**). Intriguingly, we also noted that adrenergic tumors were enriched for CD276 (B7-H3) expression, an immune checkpoint marker that protects neuroblastoma cells from NK cell-mediated cytotoxicity, which may partly account for the decreased immune response signatures in these cells^14^ (**Supplementary Fig. 4b**). We next quantified the immune response signatures in a subset of tumors (n = 100) from both the upper and lower quartiles of the M-A score that had significant differences in their activation and evasion scores (**Supplementary Fig. 4c, d**). High M-A scoring (mesenchymal) tumors had significantly higher expression levels of cytotoxic T and NK cell signatures compared with those of low M-A scoring (adrenergic) tumors (**Fig. 2f**). Considering that the M-A score and, to a lesser extent, higher *MYC* expression were positively associated with IA and IE scores in pairwise testing (**Supplementary Fig. 3f**), we next assessed their relative contributions as independent predictors of an immune response in a multivariate multiple regression model consisting of immune activation and evasion scores and T and NK cell signatures (**Fig. 2g**). Because *MYCN* amplification is linked to immune suppression, we also included the 157-gene *MYCN* signature generated by Valentijn et al^27^ and subsequently used by Wei et al to identify immune predictors in *MYCN-*nonamplified neuroblastoma^25^. In this analysis also, the M-A score was a better predictor of tumor immunogenicity than *MYC* expression or *MYCN* signature (**Fig. 2g**).

Second, we used CIBERSORT (Cell type Identification By Estimating Relative Subsets Of known RNA Transcripts)^43^, as a deconvolution approach to estimate the fraction of immune cell infiltration associated with the adrenergic and mesenchymal lineage tumors. We first established the fraction of tumor-infiltrating leukocytes (TILs) in the most mesenchymal versus the most adrenergic tumors (n = 6 each), observing significantly higher fractions in mesenchymal tumors (**Supplementary Fig. 4e**). Next, we quantified the average immune cell content in 100 tumors from the upper and lower quartiles of the M-A score (**Fig. 2h**). Consistent with our previous results (**Fig. 2f**), CIBERSORT analysis also showed significantly increased enrichment for cytotoxic CD8^+^ T cells in mesenchymal tumors, which was also associated with a concomitant increase in regulatory T cells, including those expressing markers of T-cell exhaustion (**Fig. 2h; Supplementary Fig. 4e**). Interestingly, we also observed that mesenchymal tumors comprised a higher fraction of naïve B cells, which were recently shown to take part in antitumor immunity ^44^ (**Fig. 2h; Supplementary Fig. 4e**). Together, our findings indicate that the mesenchymal cell state is a strong predictor of neuroblastoma immunogenicity.

### Tumor cell-intrinsic upregulation of immune pathways in mesenchymal neuroblastoma

We next sought to understand the extent to which the presence or absence of an immunogenic signature in the bulk RNA-sequencing data was intrinsic to tumor cells or was conferred by the tumor microenvironment. Analysis of the lineage identities of a panel of 24 human neuroblastoma cell lines (15 *MYCN-*amplified; 9 *MYCN-*nonamplified) from RNA-sequencing data (GSE28019) revealed a gradient of adrenergic-to-mesenchymal scores (**Supplementary Fig. 5a**). Consistent with our observations in primary tumors, cell lines with higher mesenchymal gene signatures grouped together and had significantly higher expression of tumor cell-intrinsic immune genes, compared with the remainder, which had higher adrenergic scores and were mostly associated with reduced immune marker gene expression (**Fig. 3a, b**). To further understand the association of tumor cell-intrinsic immune pathways with lineage state, we focused on two neuroblastoma cell lines –SH-SY5Y and SH-EP – subclones of the *MYCN-*nonamplified SK-N-SH cell line separated on the basis of neuroblastic versus substrate-adherent morphology^45^ and determined to be adrenergic and mesenchymal^38^, respectively. RNA sequencing showed that the differentially up- and down-regulated genes in SH-EP compared with SH-SY5Y cells significantly overlapped with established signatures of mesenchymal and adrenergic states, thus confirming their respective phenotypes (**Supplementary Fig. 5b, c**). We noted that genes with roles in eliciting an immune response were among the top differentially upregulated genes in mesenchymal SH-EP cells, especially those involved in antigen processing and presentation and positive regulation of MHC expression (**Fig. 3c, d**). Moreover, gene ontology (GO) analysis of the upregulated transcripts revealed enrichment for innate and adaptive immune responses including type-I interferon signaling and ligands for the NK cell receptor, NKG2D (NK cell lectin-like receptor, KLRK1) (**Supplementary Fig. 5d**). By contrast, adrenergic SH-SY5Y cells that showed upregulation of neuronal lineage markers did not show significant enrichment of immune function genes (**Fig. 3d**), providing further evidence that cell-intrinsic immunogenicity is associated with the mesenchymal phenotype.

**Fig. 3.**
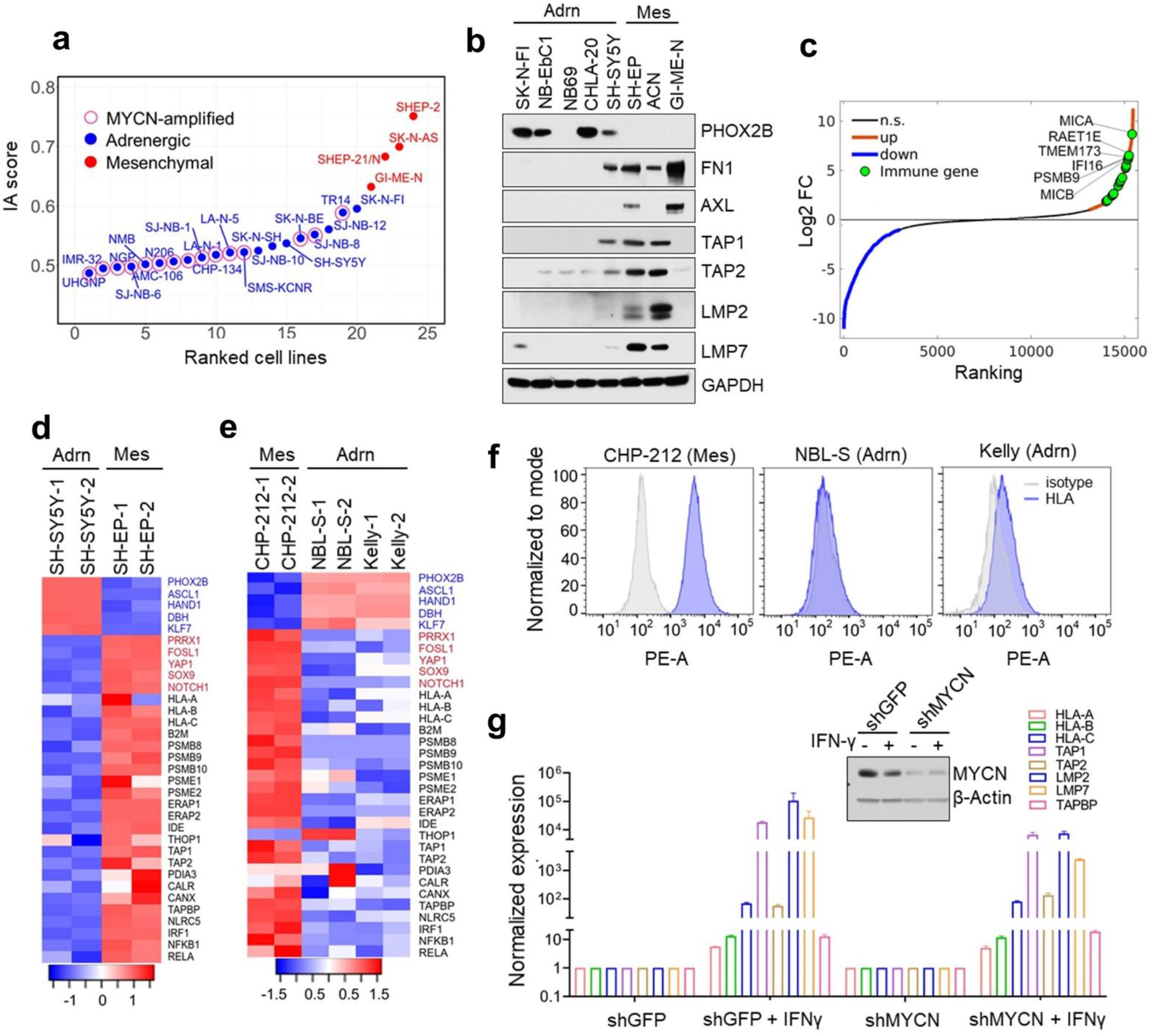
Tumor cell-intrinsic immune marker genes are upregulated in mesenchymal NBs. (**a**) Scatter plot of the immune activation (IA) scores of human neuroblastoma cell lines (RNA-seq data; GSE28019). Cell lines are arranged based on increasing IA scores and designated as adrenergic or mesenchymal based on lineage-specific gene expression. (**b**) Western blot (WB) analysis of adrenergic (PHOX2B) and mesenchymal (FN1, AXL) cell lineage markers and antigen processing genes (TAP1/2, LMP2/7) in *MYCN-*nonamplified NB cell lines. GAPDH was used as the loading control. Adrn, adrenergic; Mes, mesenchymal. (**c**) Waterfall plot of the fold-change in RNA expression levels of up- and downregulated genes in SH-EP compared to SH-SY5Y NB cells; selected immune genes are highlighted in green. (**d, e**) Heat maps of lineage marker (*blue*, adrenergic; *red*, mesenchymal) and MHC and antigen processing machinery gene (*black*) expression in the indicated *MYCN-*nonamplified (**d**) and *MYCN-*amplified (CHP-212, Kelly) and overexpressing (NBL-S) (**e**) adrenergic and mesenchymal cells (*n* = 2 biological replicates). Rows are z-scores calculated for each transcript in each cell type. (**f**) Fluorescence activated cell sorting (FACS) analysis of cell surface HLA expression in the cells depicted in **e**. Isotype controls are depicted in gray. The X-axis denotes fluorescence intensity of indicated proteins using phycoerythrin (PE-A) tagged antibodies. Results representative of 2 independent experiments. (**g**) RT-qPCR analysis of antigen processing and presentation genes in *MYCN-*amplified Kelly NB cells engineered to express shMYCN or shGFP (control) with or without IFN-γ induction (100 ng/mL for 24 hr.). Data are normalized to GAPDH and represent means ± SD, *n* = 2 biological replicates. Inset, WB analysis of MYCN in control and shMYCN cells. Actin was used as a loading control.

The absence of a productive immune response has often been described in *MYCN-* amplified neuroblastoma tumors^24,25^; indeed, the vast majority of such tumors in our cohort exhibited similar findings (**Supplementary Fig. 2a, b**). Nonetheless, based on our intriguing finding of upregulation of immune response genes in a small number of *MYCN-*amplified tumors (**Supplementary Fig. 2a, b**) that possessed mesenchymal cell signatures (**Fig. 2e**), we sought to understand the role of MYCN in mediating this immune response. We used Kelly and CHP-212 human neuroblastoma cells that expressed amplified *MYCN* but were of adrenergic and mesenchymal phenotypes, respectively^37,38^, and NBL-S cells that lacked *MYCN* amplification but expressed moderate levels of MYCN RNA and protein^46^ (**Supplementary Fig. 5e**) and were classified as adrenergic (van Groningen et al., 2017 and this study). RNA-sequencing and flow cytometry analysis suggested that tumor cell-intrinsic immune genes involved in antigen processing and MHC regulation were highly expressed in mesenchymal CHP-212 compared to adrenergic Kelly and NBL-S cells (**Fig. 3e, f; supplementary fig. 5e**). Importantly, although MYCN expression in NBL-S cells was lower than in Kelly cells (**Supplementary Fig. 5f**), these immune transcripts were expressed at lower levels in both cell lines, consistent with their adrenergic status (**Fig. 3e; Supplementary Fig. 5f**). To further verify that cell state dictate tumor cell-intrinsic immunogenicity, we depleted MYCN expression in Kelly cells and observed no significant change to the IFN-γ-induced expression of HLA and antigen processing genes compared to control cells (**Fig. 3g**). Thus, our findings suggest that the lineage state of neuroblastoma cells specifies the expression of tumor cell-intrinsic immune marker genes.

### Cellular reprogramming to the mesenchymal state leads to increased immunogenicity

We next questioned whether acquisition of the mesenchymal phenotype would be sufficient to render adrenergic neuroblastoma cells immunogenic. One of the top overexpressed genes in SH-EP mesenchymal cells, *PRRX*1, encodes a core lineage-specific homeobox transcription factor (TF), whose overexpression induces the mesenchymal state in neuroblastoma cells^37,38^ (**Supplementary Fig. 5b**). We therefore overexpressed doxycycline-inducible *PRRX1* in adrenergic SH-SY5Y cells and observed a gradual loss of the adrenergic lineage marker PHOX2B, together with increased expression of the mesenchymal markers, fibronectin, vimentin and AXL (**Fig. 4a; Supplementary Fig. 6a**). By contrast, overexpression of DNA-binding mutants of *PRRX1* that contained homeodomain deletions had no effect on mesenchymal marker expression, indicating that the lineage switch was a direct consequence of PRRX1-mediated transcriptional control (**Fig. 4a**). Next, to determine whether the phenotypic switch had any effect on tumor cell-intrinsic pro-inflammatory pathways, we analyzed the expression of genes involved in antigen processing (*TAP1, TAP2, LMP2, LMP7*) as well as *IFI16* and *STING* (*TMEM173*), innate immune regulators that were differentially upregulated in mesenchymal NB cells (**Fig. 3c**). Induction of wild-type (WT) *PRRX1* led to increased RNA expression of these genes (**Fig. 4b**). Moreover, WT PRRX1 but not its DNA-binding mutants led to increased TAP1 and LMP7 protein expression, which was accompanied by a sustained increase in cell surface MHC expression (**Fig. 4a, c**). Additionally, PRRX1 induction led to increased cell surface expression of MICA and MICB, ligands for the activating NK cell receptor NKG2D, in a minor population of cells (**Fig. 4d**), in agreement with elevated expression of these proteins in mesenchymal neuroblastoma cells (**Fig. 3c**). These results suggest that conversion from the adrenergic to mesenchymal cell state may be adequate to reprogram immune-insensitive cells toward immunocompetency.

**Fig. 4.**
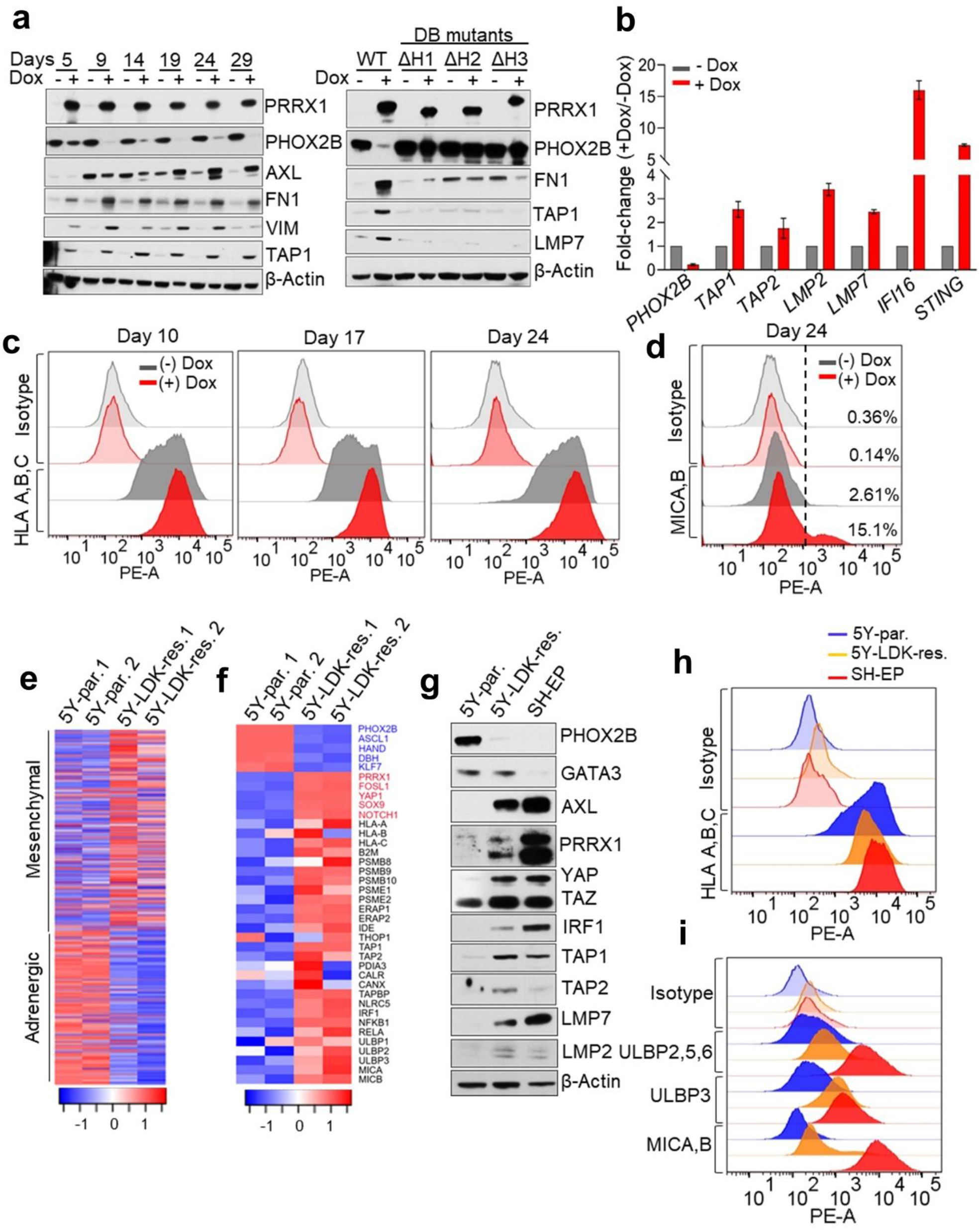
Reprogramming of adrenergic NB cells to the mesenchymal cell state leads to increased expression of immune response genes. **(a)** *Left*, WB analysis of PHOX2B and antigen processing gene expression in adrenergic SH-SY5Y cells engineered to express doxycycline (dox)-inducible PRRX1 in the presence or absence of dox (200 ng/mL) at the indicated time points. *Right*, WB analysis of the indicated proteins in SH-SY5Y cells expressing dox-inducible wild-type (WT) or DNA-binding mutants of PRRX1 at 10 days post dox-induction. The DNA-binding (DB) mutants harbor individual deletions of the three α-helices (ΔH1, ΔH2 and ΔH3) within the PRRX1 homeodomain. (**b**) RT-qPCR analysis of the indicated immune response genes in the same cells as in (**a**). Data represent the means ± SD, *n* = 2 biological replicates. (**c**) FACS analysis of cell surface HLA expression following dox-inducible expression of PRRX1 in SH-SY5Y cells at the indicated time points. Data are representative of 2-3 independent experiments. (**d**) FACS analysis of cell surface MICA/MICB expression after PRRX1 induction for 24 days in the same cells as in (**c**). A logscale expression value of 10^3^ was used as a threshold (vertical line) to gate MICA/MICB negative (<10^3^) and positive (≥10^3^) populations. Numbers on the right indicate the percentage of MICA/MICB-positive cells. Plots are representative of 2 independent experiments. (**e**) Heat map representation of adrenergic and mesenchymal gene signatures in parental (5Y-par) and LDK-resistant (5Y-LDK-res) SH-SY5Y cells (*n* = 2 biological replicates). Rows represent z-scores of log_2_ expression values for each gene in both cell types. (**f**) Heat map depicting the expression of cell lineage markers (blue, adrenergic; red, mesenchymal), antigen processing machinery genes and NKG2D ligands (black) in parental and LDK-resistant SH-SY5Y cells (*n* = 2 biological replicates). Rows represent z-scores of log_2_ expression values. (**g**) WB analysis of lineage marker and antigen processing gene expression in the indicated cells. Actin was used as a loading control in all immunoblots. (**h, i**) FACS analysis of cell surface HLA (**h**) and NKG2D ligand (**i**) expression in the indicated cells.

Transition from the adrenergic to the mesenchymal state in neuroblastoma is accompanied by resistance to chemotherapy ^38^. Whether this transition in the face of treatment pressure might include the acquisition of a pro-inflammatory signature is unclear, leading us to compare adrenergic neuroblastoma cells that had gained mesenchymal features during the development of treatment resistance with their sensitive, adrenergic counterparts. For this purpose, we used an isogenic pair of cell lines comprising adrenergic SH-SY5Y neuroblastoma cells that express the *ALK^F1174L^* mutation and are sensitive to the small molecule inhibitor ceritinib (LDK378) (parental SH-SY5Y, IC_50_ = 150 nM), and their ceritinib-resistant derivatives (LDK-resistant SH-SY5Y, IC_50_ = 1101 nM) (**Supplementary Fig. 6b**)^47^. Comparison of the gene expression signatures of these cell lines revealed significant downregulation of adrenergic transcripts in LDK-resistant SH-SY5Y cells with concomitant upregulation of the mesenchymal signature (**Fig. 4e**). Moreover, a significant overlap was noted between the differentially up- or downregulated transcripts in the LDK-resistant SH-SY5Y cells and established signatures of mesenchymal and adrenergic states, respectively (**Supplementary Fig. 6c**), suggesting that these cells had acquired features of the mesenchymal phenotype with resistance. Consistent with the key role of PRRX1 in triggering the conversion from an adrenergic to mesenchymal cell state, we observed that this TF was among the top upregulated genes in LDK-resistant SH-SY5Y cells (**Supplementary Fig. 6d**). The mesenchymal state of the LDK-resistant SH-SY5Y cells was further supported by the loss of the pivotal adrenergic marker, PHOX2B, and increased expression of additional mesenchymal markers AXL, YAP, TAZ, and IRF1, although these changes were not as pronounced as those in SH-EP mesenchymal cells that served as a positive control (**Fig. 4f, g**). Further evidence supporting the conversion to the mesenchymal state came from the differential upregulation in LDK-resistant SH-SY5Y cells of cell-intrinsic immune markers engaged in antigen processing and presentation and NK cell activating receptor ligands (PSMB9, MICA, MICB); in fact, these were among the top differentially upregulated genes in LDK-resistant SH-SY5Y cells (**Supplementary Fig. 6d, e; Fig. 4f, g**). These changes in immune genes coincided with increases in cell surface expression of MHC receptors to levels comparable to those in mesenchymal SH-EP cells (**Fig. 4h**), as well as the increased expression of ligands for the NK cell-activating receptor NKG2D (**Fig. 4i**). Thus, the genetic reprogramming from the adrenergic to the mesenchymal state that occurred with therapy resistance also led to the upregulation of tumor cell-intrinsic pro-inflammatory pathway genes suggesting that such conversion could render the tumor cells susceptible to recognition by T and NK cells.

### Immune response gene expression during cell state transition is epigenetically regulated

As lineage plasticity in neuroblastoma is epigenetically driven^37,38,48,49^, we next questioned whether the altered expression of immune response genes observed in the individual cell states could be the result of changes in chromatin organization. To this end, we analyzed the chromatin occupancies of active and repressive histone marks at immune genes that were upregulated in adrenergic SH-SY5Y cells upon induction of PRRX1 (**Fig. 4b**). Indeed, PRRX1 induction resulted in increased binding of the active H3K4me3 mark as well as loss of repressive H3K27me3 binding at several candidate immune genes, including the APM genes *TAP1* and *PSMB9* (**Fig. 5a; Supplementary Fig. 7a**). To understand epigenetic modifications that occur during the spontaneous transition between the two lineage states (as compared with forced expression of *PRRXI*) on a genome-wide basis, we compared histone occupancies between adrenergic (parental SH-SY5Y) cells and those that had acquired mesenchymal characteristics with drug resistance (LDK-resistant SH-SY5Y) (**Fig. 4e**), using SH-EP cells as a typical example of the mesenchymal state. ChIP-seq analysis of active H3K27ac binding identified that the super-enhancers (SEs) in LDK-resistant SH-SY5Y cells were associated with genes that conferred mesenchymal identity while parental SH-SY5Y cells retained SEs at genes that conferred adrenergic identity (**Supplementary Fig. 7b**), consistent with evidence that lineage plasticity is driven by cell type-specific SEs^37,38^. To determine whether the SE-mediated regulation of lineage genes also extended to genes associated with immune responsiveness, we analyzed the genes in our 41-gene immune activation signature (**Supplementary Table 3; Fig. 1e**) as well as those associated with an IFN-response signature (n = 91) in primary tumors and cell lines (**Fig. 1b; Supplementary Fig. 5d**, see Methods). Despite the higher expression of these genes in mesenchymal cells (LDK-resistant SH-SY5Y and SH-EP), none was associated with an SE, prompting us to focus on the promoter regions. We observed significantly higher enrichment of H3K27ac and H3K4me3 binding at regions spanning the transcription start sites (TSS ± 2 kb) at cell-intrinsic immune genes in LDK-resistant SH-SY5Y and SHEP compared to parental SH-SY5Y cells (**Fig. 5b, c**). On the other hand, the adrenergic parental SH-SY5Y cells showed significantly higher occupancies of the H3K27me3 repressor mark at these immune gene promoters (**Fig. 5b, c**). Analysis of the polycomb repressive complex 2 (PRC2) that promotes H3K27me3 deposition at repressed chromatin^50^ revealed that immune response genes enriched for H3K27me3 binding, such as those encoding the NKG2D ligands *MICA/B*, *ULBP2/3* and *RAET1G* had significantly higher occupancies for PRC2 subunits, EZH2 and SUZ12 in adrenergic parental SH-SY5Y cells compared to negative control regions that lacked H3K27me3 binding (*PHOX2B* and *LIN28B*) (**Supplementary Fig. 7c**), suggesting active immune gene repression in these cells.

**Fig. 5.**
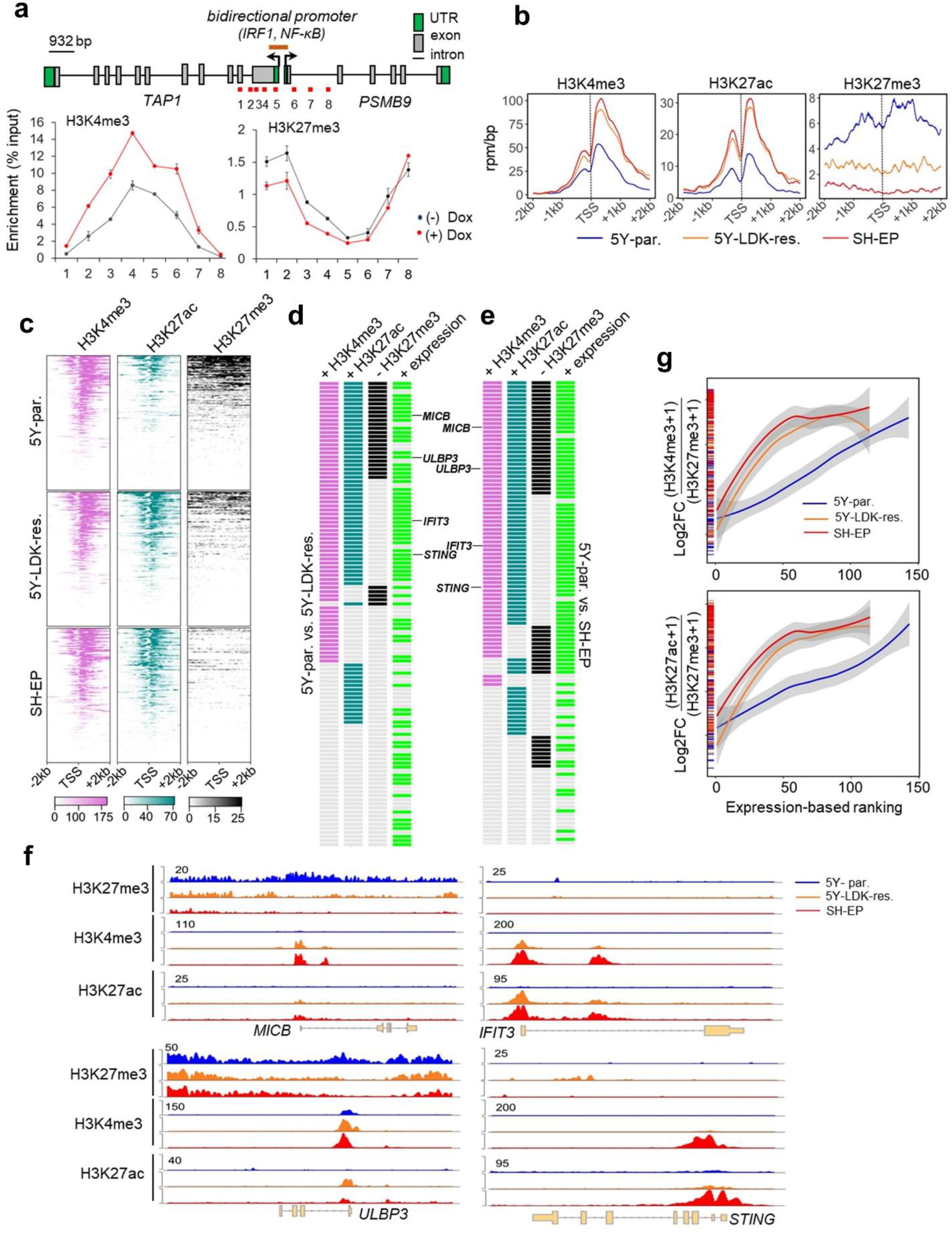
Activation of immune gene expression associated with cell state transition is epigenetically regulated. (**a**) *Upper*, Linear representation of TAP1 and PSMB9 gene loci showing the locations of the bidirectional promoter and IRF1 and NF-κB binding sites. The amplicons (1-8) analyzed for histone mark occupancy are shown in red. *Lower*, ChIP-qPCR analysis of H3K4me3 and H3K27me3 enrichment at the indicated amplicons along the TAP1/PSMB9 locus in adrenergic SH-SY5Y cells expressing dox-inducible PRRX1 in the presence or absence of dox (200 ng/mL) for 10 days. Data represent the means ± SD, *n* = 2 biological replicates. (**b**) Metagene representations of average ChIP-seq occupancies of the indicated histone marks at the promoters of tumor cell-intrinsic immune response genes (TSS ± 2 kb; n = 134) in parental (5Y-par.), LDK-resistant (5Y-LDK-res.) SH-SY5Y and SH-EP NB cells. (**c**) Heat map representation of histone enrichment at the same immune gene promoters as in (**b**), ranked in decreasing order of occupancy in the indicated cells. Each row represents the normalized densities of histone marks within a ± 2 kb window centered on the TSS. (**d, e**) Representation of pairwise comparisons between parental SH-SY5Y and LDK-resistant SH-SY5Y (**d**), and parental SH-SY5Y and SH-EP cells (**e**). The changes (+, gained; -, lost) in occupancies of the active (H3K4me3, H3K27ac) and repressive (H3K27me3) histone marks (log_2_ FC ≥ 0.75, TSS ± 2 kb), together with the corresponding changes in RNA expression (+, overexpressed; log_2_ FC ≥ 1) of each of the 134 tumor cell-intrinsic immune genes analyzed in (**b**) are shown. Representative genes showing either a switch from repressive to active chromatin (*MICB, ULBP3*) or associated only with a gain of active chromatin (*IFIT3, STING*) are shown. (**f**) ChIP-seq tracks depicting the gain of active histone binding together with the loss of repressive histone binding *(left)* or gain of active marks without changes in repressive mark occupancy *(right)* at the indicated immune gene loci. Signal intensity is given at the top left corner for each track. (**g**) Loess regression analysis of the correlation between the ratios of active to repressive histone binding at the promoters (TSS ± 2kb) of immune response genes and their RNA expression (*Upper*, H3K4me3:H3K27me3; *Lower*, H3K27ac:H3K27me3). Genes are ranked based on increasing expression. Shaded regions represent 95% confidence intervals.

We next sought to understand whether the activation of immune response genes observed during the cell state transition from sensitivity to resistance represented a *switch* from repressive to active chromatin or a *gain* of active chromatin marks. To this end, we quantified the changes in histone binding occupancies between adrenergic parental SH-SY5Y and mesenchymal LDK-resistant SH-SY5Y or SH-EP cells using pair-wise comparisons (**Supplementary Fig. 7d, e**). Compared to parental SH-SY5Y, LDK-resistant SH-SY5Y and SH-EP cells gained significant H3K4me3 binding at the promoters of 60% and 62% immune genes (68 and 71 of 114) respectively, which corresponded with their increased expression (**Fig. 5d, e; Supplementary Fig. 7f**). A similar significant enrichment of the H3K27ac histone mark was observed at the promoters of these immune genes [LDK-resistant SH-SY5Y, 58% (66/114); SH-EP, 67% (76/114)]. Interestingly, gain of these active marks was accompanied by a concomitant loss of H3K27me3 repressive histone binding at the promoters of 25% and 35% of (29 and 40 of 114) immune genes in LDK-resistant SH-SY5Y and SH-EP cells respectively, as represented by the NKG2D ligands, *MICB* and *ULBP3* (**Fig. 5d-f; Supplementary Fig. 7f**). On the other hand, in LDK-resistant SH-SY5Y and SH-EP mesenchymal cells, 48% and 41% (55 and 47 of 114) immune genes, such as the IFN-regulated factors *IFIT3 and STING*, gained either one or both active marks without changes in occupancy of the repressive mark (**Fig. 5d-f; Supplementary Fig. 7f**). Furthermore, the mesenchymal cells showed a significantly higher ratio of active to repressor histone binding at the TSSs of immune-related genes (H3K4me3 or H3K27ac/H3K27me3) (**Supplementary Fig. 7g**), which importantly, also correlated with the increased expression of these genes in this cell state (**Fig. 5g**). Therefore, our results suggest that the immune gene activation observed with the transition from the adrenergic to the mesenchymal cell state represents either a switch from repressive to active chromatin or a gain of active chromatin at promoter regions.

### Mesenchymal neuroblastoma cells functionally engage cytotoxic T cells

To assess the functional consequences of the increased immunogenicity associated with a mesenchymal phenotype, we utilized the murine neuroblastoma cell line NB-9464, which was derived from tumors arising in the Th-MYCN genetically engineered mouse model (GEMM). This model was generated in immunocompetent C57BL/6 mice and the tumors recapitulate the genetic and immunological features of human neuroblastoma^51,52^. We observed that NB-9464 cells consisted of distinct populations that could be sorted on the basis of surface MHC class I H-2Kb expression into high (H-2Kb^hi^)- or low (H-2Kb^lo^)-expressing populations (**Supplementary Fig. 8a; Fig. 6a**), both of which expressed transgenic human MYCN (**Fig. 6b**). Consistent with our hypothesis, H-2Kb^hi^ cells were enriched for *bona fide* mesenchymal markers - Prrx1, Sox9, Notch1, and Snai2 (Slug) (**Fig. 6b**), and showed enhanced migration and invasion, as might be expected from their neural crest cell-like state (**Supplementary Fig. 8b, c**). Importantly, the mesenchymal H-2Kb^hi^ NB-9464 cells, but not their adrenergic H-2Kb^lo^ counterparts showed augmented expression of cell surface class I MHC H-2Kb in response to IFN-γ, suggesting that the mesenchymal cells had the potential for inducing a T cell-driven antitumor immune response (**Fig. 6c**).

**Fig. 6.**
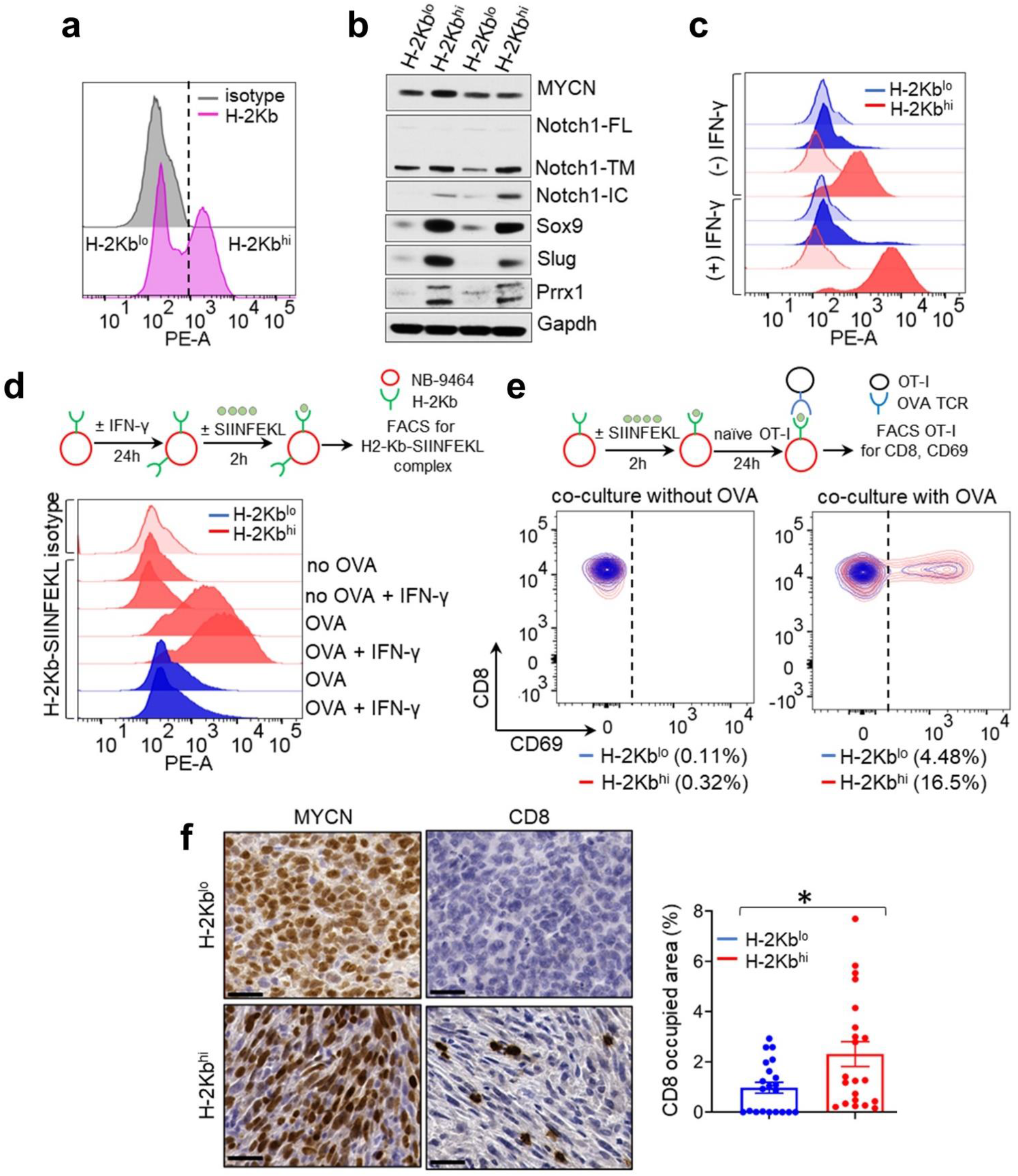
Mesenchymal NB cells functionally engage cytotoxic T cells. (**a**) FACS analysis of H-2Kb expression in unsorted NB-9464 cells. Vertical dashed line denotes the logscale expression value used as a threshold to gate H-2Kb^lo^ and H-2Kb^hi^ cell populations. (**b**) WB analysis of the indicated lineage markers in H-2Kb^lo^ and H-2Kb^hi^ cell populations. Notch1-FL, full length; -TM, transmembrane; -IC, intracellular. GAPDH is used as a loading control. (**c**) FACS analyses of basal and IFN-γ-induced (100 ng/mL for 24 hr.) surface H-2Kb expression (darker colored histograms) in H-2Kb^lo^ and H-2Kb^hi^ cells compared to isotype controls (lighter colored histograms). Plots representative of 2 independent experiments. (**d**) *Upper*, Schematic of OVA binding assay. *Lower*, FACS analysis of surface H-2Kb-bound SIINFEKL OVA peptide in H-2Kb^hi^ and H-2Kb^lo^ cells under basal or IFN-γ-induced conditions as in (**c**) and in the absence or presence of the OVA peptide. Plots representative of 2 independent experiments. (**e**) *Upper*, Schematic of NB-9464-OT-I co-culture assay. *Lower*, Contour plots showing the percentage of naïve OT-I cells that were activated (CD8^+^ CD69^+^) following co-culture with H-2Kb^lo^ and H-2Kb^hi^ cells for 24 hr. with or without the OVA peptide. OT-I activation was measured by FACS analysis of cell surface CD69. (**f**) *Left*, Immunohistochemical (IHC) staining for MYCN and CD8 expression in representative murine NB xenograft tumors derived from NB-9464 H-2Kb^lo^ (adrenergic) and H-2Kb^hi^ (mesenchymal) cells in immunocompetent syngeneic (C57BL/6) mice. Scale bars, 100 µm. *Right*, Bar graphs showing the percentage of area occupied by CD8^+^ T cells in H-2Kb^lo^ (0.9% ± 0.2%) vs. H-2Kb^hi^ (2.3% ± 0.5%) tumors; *\*P* < 0.05, two-tailed Welch’s t-test. Each dot represents one of three independent measurements for each tumor. Data represent mean ± SEM.

Hence, to determine whether the increased MHC class I expression of mesenchymal versus adrenergic cells translated into T cell activation, we first asked whether mesenchymal H-2Kb^hi^ NB-9464 cells were capable of exogenous antigen presentation. Using the well-characterized chicken ovalbumin-derived peptide (OVA_257-264_ or SIINFEKL) antigen that binds to H-2Kb and can be recognized by specific T cell receptors (TCRs) on CD8^+^ T cells^53^, we found that in comparison to H-2Kb^lo^ cells, H-2Kb^hi^ cells expressed significantly higher levels of the H-2Kb-SIINFEKL complex (**Fig. 6d**). Next, we determined whether antigen presentation through H-2Kb enables mesenchymal tumor cells to be recognized by antigen-specific T cell receptors (TCRs) on CD8^+^ T cells, the first step towards a cytotoxic response. For this purpose, we used OT-I CD8^+^ T cells from C57BL/6 mice expressing a transgenic TCR that specifically recognizes the H-2Kb-SIINFEKL complex^53^. H-2Kb^hi^ or H-2Kb^lo^ cells loaded with the SIINFEKL peptide were cocultured with naïve OT-I cells, after which OT-I activation was measured through cell surface CD69 expression, an early marker of T-cell activation^54^. Mesenchymal H-2Kb^hi^ cells led to significantly higher OT-I activation in comparison with adrenergic H-2Kb^lo^ cells, indicating specific recognition of the H-2Kb-SIINFEKL complex by the TCR on OT-I cells (**Fig. 6e**). By contrast, co-cultures of OT-I cells and either H-2Kb^lo^ or H-2Kb^hi^ NB-9464 neuroblastoma cells without the SIINFEKL peptide did not lead to T-cell recognition, confirming the specificity of the TCR-antigen interaction (**Fig. 6e**). Finally, we investigated whether the differential MHC-I expression between adrenergic and mesenchymal neuroblastoma cells influenced tumor growth *in vivo* through subcutaneous injection of H-2Kb^hi^ or H-2Kb^lo^ cells into syngeneic C57BL/6 (H-2Kb haplotype) mice (**Supplementary Fig. 8d**). We noted an earlier onset of tumor formation with H-2Kb^lo^ cells compared to H-2Kb^hi^ cells (**Supplementary Fig. 8d**). However, once consistent tumor growth was established, growth or survival rates did not change substantially between the two groups (**Supplementary Fig. 8e**), despite the persistence of higher H-2Kb and Prrx1 expression in the H-2Kb^hi^ tumors compared with the H-2Kb^lo^ tumors (**Supplementary Fig. 8f**,**g**). While both types of tumor cells had MYCN expression, histologically, in keeping with their adrenergic phenotype H-2Kb^lo^ tumors were composed of densely arranged small round blue cells, whereas H-2Kb^hi^ tumors predominantly comprised elongated, spindle-like cells interspersed with clusters of small round blue cells (**Fig. 6f**). We next analyzed the immune status of these tumors, reasoning that tumors arising from immunogenic H-2Kb^hi^ cells would be infiltrated by T cells. Indeed, H-2Kb^hi^ tumors showed significantly higher CD8^+^ T cell infiltration compared with H-2Kb^lo^ tumors (**Fig. 6f**). Taken together, these results suggest that the immunogenic traits of mesenchymal neuroblastoma cells translate into the recruitment of cytotoxic T cells into the tumor microenvironment.

### Mesenchymal neuroblastoma cells induce NK cell degranulation

We next analyzed the functional relevance of the increased expression of the NKG2D NK cell receptor ligands (ULBP1-3, MICA, MICB) seen in mesenchymal neuroblastoma tumors (**Supplementary Fig. 4c; Fig. 4i**). Since the interaction of the receptor with its cognate ligands on target cells is the first step towards a cytotoxic response (**Fig. 7a**), we compared the ability of adrenergic (parental SH-SY5Y) and mesenchymal (LDK-resistant SH-SY5Y and SH-EP) cells to bind to the recombinant NKG2D receptor fusion protein in an *in vitro* binding assay. Both mesenchymal cell types showed increased binding to the NKG2D receptor, in keeping with their increased cell surface expression of the ULBP2/3/5/6, MICA and MICB ligands (**Fig. 7b**). NK cell cytotoxicity is mediated by granzyme proteases and the pore-forming protein perforin, which in resting cells are stored in secretory lysosomes or lytic granules marked by lysosome-associated membrane protein-1 (LAMP-1 or CD107a)^55^. Upon target recognition, NK cells undergo degranulation, or exocytosis of the lytic granules, which is associated with relocation of the CD107a antigen to the cell membrane. Using cell-surface CD107a as a specific marker of degranulation, we measured degranulation of peripheral blood NK cells harvested from healthy human donors in the presence of parental SH-SY5Y, LDK-resistant SH-SY5Y and SH-EP targets.

**Fig. 7.**
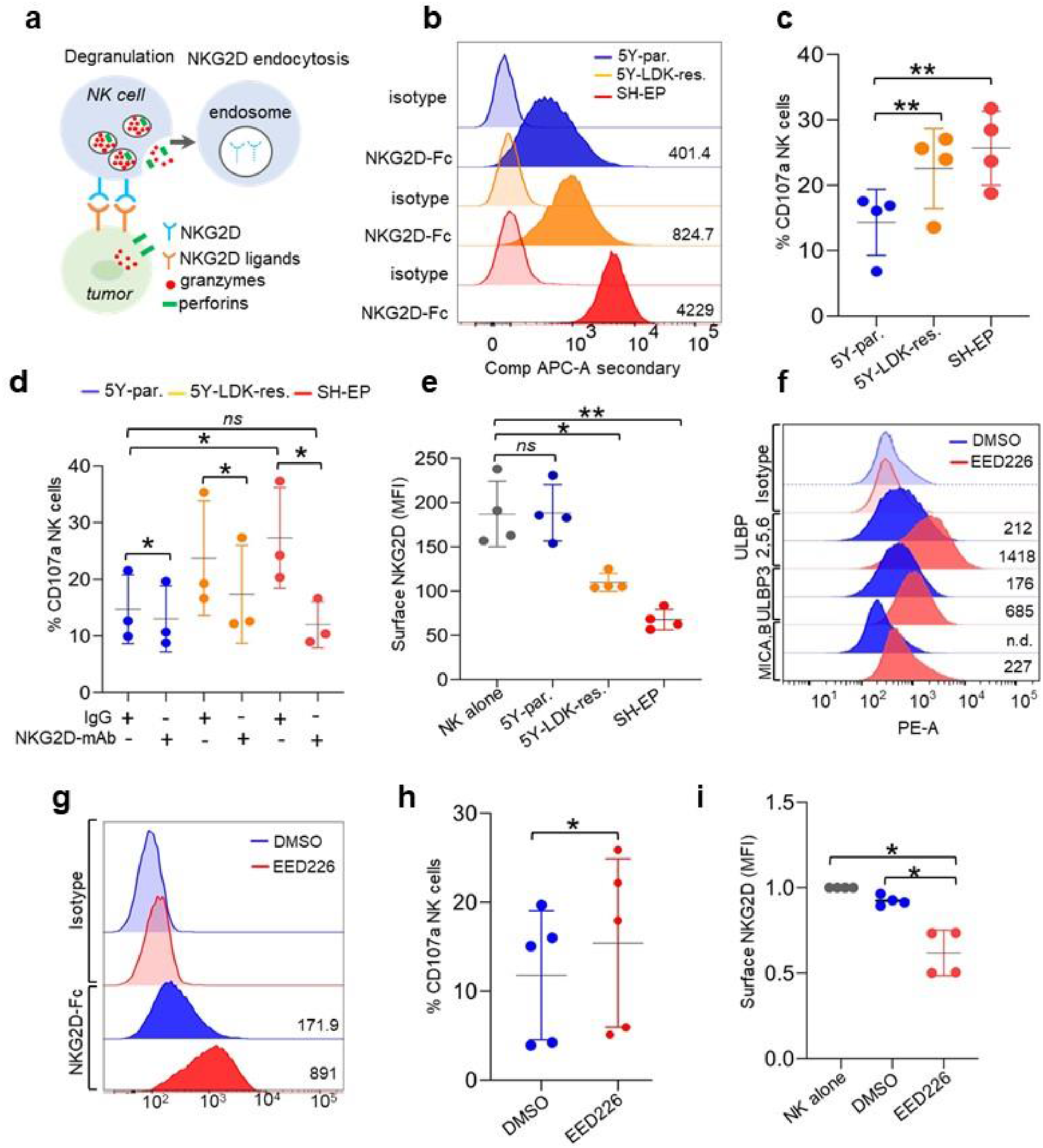
Mesenchymal NB cells induce NK cell degranulation. (**a**) Schematic representation of the interaction between NKG2D receptors on NK cells and cognate ligands on tumor cells, leading to NK cell degranulation and receptor endocytosis. (**b**) FACS analysis of purified human NKG2D-Fc protein binding (darker histograms) to adrenergic parental SH-SY5Y, and mesenchymal LDK-resistant SH-SY5Y and SH-EP cells. Comp Alexa-647, compensated fluorescence intensity of NKG2D-Fc protein detected using Alexa 647-conjugated anti-human IgG. Lighter histograms indicate staining with Alexa 647-conjugated anti-human IgG only. Numbers indicate median fluorescence intensity (MFI). Plots representative of 2 independent experiments. (**c**) X-Y plot showing the percentage of degranulating NK cells following co-culture with the same cells as in (**b**) for 4 hr. at an effector: target (E: T) cell ratio of 1:2. Degranulation was measured by FACS analysis of cell-surface CD107a. Data represent the means ± SD, *n* = 4 biological replicates. (**d**) X-Y plot showing the effect of a control IgG1 or an NKG2D blocking antibody on NK cell degranulation following co-culture with the indicated cells for 4 hr. NK cell degranulation was measured as in (**c**). Data represent the means ± SD, *n* = 4 biological replicates. (**e**) X-Y plot depicting the MFI of NKG2D expression measured by FACS on naïve NK cells (NK alone) or following co-culture with parental SH-SY5Y, LDK-resistant SH-SY5Y and SH-EP cells for 4 h. Data represent the means ± SD, *n* = 4 biological replicates. (**f**) FACS analysis of surface ULBP2/5/6, ULBP3, and MICA/MICB in adrenergic parental SH-SY5Y cells treated with DMSO (vehicle control) or EED226 (5 µM for 8 days). (**g**) FACS analysis of purified human NKG2D-Fc protein binding to parental SH-SY5Y cells treated with either DMSO or EED226 as in (**f**). Light gray histograms indicate human IgG1 isotype control. Numbers on the right represent MFI values for FACS plots. Plots in (**f**) and (**g**) are representative of 2 independent experiments. (**h, i**) X-Y plots showing NK cell degranulation (**h**) and MFI of surface-NKG2D (**i**) following co-culture of naïve NK cells with parental SH-SY5Y cells treated with DMSO or EED226 as in (**f**). NK cell granulation and NKG2D MFI were measured as in (**c**) and (**e**). Data represent means ± SD, *n* = 4 (**h**) and 5 (**i**) biological replicates. Significance for all results was calculated using the paired two-tailed Student’s t-test (\**P* < 0.05; \*\**P* < 0.01); *ns*, not significant

Co-culture of NK cells with LDK-resistant SH-SY5Y and SH-EP cells, both of which express ligands for the NKG2D receptor, resulted in increased NK cell degranulation compared to parental SH-SY5Y cells that did not express these ligands (**Fig. 7c**). To confirm that the increased NK cell degranulation in LDK-resistant SH-SY5Y and SH-EP cells was specific to the NKG2D receptor, we blocked its function with an anti-NKG2D monoclonal antibody. Compared to the isotype control, blockade of NKG2D receptor activity completely abrogated the increased NK cell degranulation in LDK-resistant SH-SY5Y and SH-EP cells, but had no effects on parental SH-SY5Y cells, signifying that the modest but robust increase in degranulation in the presence of mesenchymal cells was specific to the NKG2D receptor on NK cells (**Fig. 7d**). Upon interaction with their cognate ligands on target cells, NKG2D receptors are internalized via ubiquitin-dependent endocytosis leading to their lysosomal degradation, rendering the loss of surface NKG2D receptor expression a robust readout for ligand-receptor engagement^56^ (**Fig. 7a**). In line with the presence of functional NKG2D ligands on mesenchymal cells, co-cultures with LDK-resistant SH-SY5Y and SH-EP cells led to significant downregulation of NK cell surface-associated NKG2D expression, whereas co-culture with the adrenergic parental SH-SY5Y cells did not alter the abundance of surface NKG2D expression (**Fig. 7e**). Moreover, consistent with the increased levels of NK degranulation upon co-culture with mesenchymal cells, LDK-resistant SH-SY5Y were more susceptible to NK-induced cell death compared to parental SH-SY5Y cells (**Supplementary Fig. 9a**)

The observation that genes encoding NKG2D ligands are repressed by the PRC2 complex in adrenergic parental SH-SY5Y cells (**Supplementary Fig. 7c**) prompted us to examine whether PRC2 inhibitors could induce the expression of these transcripts and influence NK cell function. Indeed, treatment of parental SH-SY5Y cells with EED226, an allosteric inhibitor of the PRC2 complex^57^, led to increased expression and surface localization of ULBP2/3 and MICA/B NKG2D ligands (**Fig. 7f**). Moreover, such increased ligand expression led to their increased binding to the NKG2D receptor fusion protein in EED226-treated cells compared to cells treated with DMSO alone (**Fig. 7g**). Consequently, PRC2 inhibition resulted in an ∼20% increase in NK cell degranulation (**Fig. 7h**). Finally, co-culture of primary NK cells with adrenergic parental SH-SY5Y cells treated with EED226 led to a significant loss of surface NKG2D receptor expression (**Fig. 7i**), suggesting that the increased degranulation resulting from PRC2 inhibition was driven by the NKG2D receptor. Overall, these results suggest that the lineage-specific expression of NK cell ligands in mesenchymal neuroblastoma cells has a functional impact on NK cell activity, and that pretreatment of adrenergic neuroblastoma cells could potentially render these cells susceptible to NK-cell mediated immunotherapy by upregulating ligand expression.

## DISCUSSION

Despite the relatively poor track record of immunotherapy for neuroblastoma, growing evidence suggests that subsets of these tumors have the potential to induce a productive immune response^9^. Here, we demonstrate that the mesenchymal cell state, characterized by neural crest cell (NCC)-like phenotypes, is a strong predictor of an antitumor immune response in neuroblastoma. Induction of this state was accompanied by the expression of tumor cell-intrinsic immune response-inducing genes that were epigenetically repressed in the more differentiated (adrenergic) tumor cells. Importantly, inhibition of the PRC2 complex relieved such repression of ligands for the activating NK cell receptor NKG2D, and led to NK cell degranulation, suggesting that this strategy could be explored as a potential measure to improve the response of patients with adrenergic neuroblastomas to NK cell-mediated therapy.

We sought to identify neuroblastomas capable of eliciting an immune response as those characterized by the differential expression of immune gene signatures while remaining agnostic to any of the established parameters that predict disease aggressiveness. Using UMAP dimension reduction to analyze gene expression data from 498 primary neuroblastoma tumors, we identified four clusters that were separated on the basis of differential activation of gene networks that regulate the antitumor immune response, neuronal differentiation, MYCN-driven processes and lipid metabolism. Intriguingly, the immunogenic cluster comprised almost equal proportions of high- and low-risk tumors, raising the possibility that the molecular mechanisms underlying immunogenicity in neuroblastoma are independent of disease aggressiveness. Thus, our analysis using a cluster-based approach enabled the identification of antitumor immune signatures as shared transcriptional programs between high- and low-risk neuroblastoma tumors, a bridging feature that would have been missed in studies based solely on differential gene expression between prognostically distinct groups of tumors.

A major finding of our study is the intimate link between tumor cell lineage and the propensity of eliciting an immune response. Neuroblastoma tumors show lineage plasticity, underscored by a phenotypic switch between undifferentiated NCC-like and more differentiated cells, two divergent cell states driven by distinct transcriptional programs ^37,38^. Our analysis revealed that neuroblastomas enriched in NCC-derived signatures showed significantly higher antitumor immunity featuring T and NK cells compared to tumors enriched for signatures of the adrenergic lineage. Our finding is substantiated by studies showing that diverse cellular states such as stemness, senescence and metastasis strongly influence the engagement of innate and adaptive immune pathways ^58–60^. Importantly, while the NCC-like/mesenchymal state promoted immune response mechanisms such as upregulation of MHC class I and infiltration of cytotoxic lymphocytes, these tumors were also characterized by the activation of immune checkpoints such as regulatory T cells and exhaustion markers linked to immune suppression, similar to those observed in chronic virally infected states ^61^. The presence of such seemingly contradictory gene signatures has important therapeutic implications for selecting patients who are likely to benefit from T cell-based immunotherapies, as agents that target negative regulatory immune checkpoints are likely to be most effective in those with a pre-existing but dampened antitumor immune response ^35^. Moreover, inhibition of cancer cell-intrinsic transcriptional programs that promote T cell exclusion have been shown to effect changes in cell state and could potentially employed to sensitize tumors to immunotherapy (Jerby-Arnon et al., 2018; Koyama et al., 2016; Spranger et al., 2015).

In contrast to other *MYC*-driven cancers where high *MYC* levels restrain inflammatory signaling and anti-tumor immune pathways^41,62^, tumors within our immunogenic cluster, enriched largely for mesenchymal/NCC signatures, had relatively high MYC expression that positively correlated with immune cell infiltration. These diametrically opposite roles of MYC in regulating immune response suggest that its transcriptional functions are likely to be very different in cancers in which aberrant MYC expression is the main oncogenic driver compared to *MYCN-*nonamplified neuroblastomas that are not dependent on MYC overexpression. This notion is supported by the observation that inhibiting endogenous MYC function in non MYC-driven pancreatic cancer models leads to decreased recruitment and retention of inflammatory cells^63^. As MYC is an essential TF in NCCs^64,65^, it is plausible that MYC promotes an immunogenic state by driving a NCC-specific transcriptional program.

We also extended the analysis between cell state and immunogenicity to *MYCN-*amplified tumors and identified a subset that incorporates both mesenchymal and immunogenic features. This finding was substantiated by our data showing that expression of functional MHC class I is retained in some *MYCN-*amplified mesenchymal neuroblastoma cells and is not perturbed by changes in MYCN levels. Indeed, a robust antitumor immune response is observed during the early stages of tumor development in Th-MYCN mice and in mouse-human chimeric tumors derived from human NCCs expressing *MYCN* and oncogenic *ALK*^17^. Considering that these mouse models are driven by gain of *MYCN* (∼4-8 copies of the *MYCN* transgene)^17,52^ rather than the amplification seen in human tumors, and that absolute levels of MYCN protein dictate transcriptional output^66^, the relatively low MYCN dosage in these tumors may account for their immunogenicity. These findings, if confirmed in additional data sets, should encourage us to reconsider the notion that all *MYCN-*amplified tumors are intrinsically immune tolerant and that a subset may in fact, be capable of inducing an immune response by virtue of their cell state.

The mutual exclusivity of the neuronal and immunogenic clusters suggests that neuroblastomas with characteristics of neuronal differentiation (i.e. adrenergic phenotype) are weakly immunogenic and thus incapable of inducing an effective immune response. Moreover, the presence of NCC-derived gene expression signatures among immunogenic tumors indicates that phenotypic reversal of sympathetic neuronal cells to an NCC-like state could contribute significantly to the antitumor immune response. Such cell state-dependent immunogenic switching could be mediated by the lineage-specific core regulatory circuitry (CRC) that drives distinct transcriptomic states in neuroblastoma. Indeed, several TFs that constitute the NCC-like (mesenchymal) CRC, including interferon regulatory factors 1-3 and IFI16, function as major drivers of tumor cell-intrinsic innate and adaptive immune responses^67,68^. Furthermore, our data identify PRRX1, another component of the mesenchymal CRC, as a regulator of MHC class I and antigen-processing gene expression in neuroblastoma. Interestingly, PRRX1 was not identified as a candidate regulator of MHC-I expression in genome-wide CRISPR knock-out screens to identify NF-κB-dependent MHC-I suppressors in neuroblastoma^69^, suggesting that the mechanisms employed by PRRX1 could be independent of NF-kB activation and may involve direct transcriptional activation of these genes. In addition, the upregulation of DNA damage sensor proteins such as IFI16 and STING in mesenchymal neuroblastomas could also contribute to the increased tumor-infiltrating lymphocyte abundance in these tumors, as suggested by results in small cell lung cancer^70^.

We have also established that the changes in immune gene expression accompanying the adrenergic to mesenchymal transition are epigenetically regulated. Unlike lineage identity genes that are regulated by super-enhancers, tumor cell-intrinsic immune genes involved in diverse immune functions such as the inflammatory response, IFN-γ signaling and NK cell recognition, are governed through changes in promoter structure, achieved by either *de novo* acquisition of permissive chromatin or an epigenetic switch from PRC2-mediated repression to a permissive chromatin landscape. Our findings support a role for the PRC2 complex in repressing genes that encode ligands for the activating NK cell receptor NKG2D in adrenergic neuroblastoma cells and the use of PRC2 inhibitors to augment the NK cell response against these cells. This approach is justified on several grounds: our results add to the growing body of evidence for tumor-cell autonomous function of PRC2 as a barrier to anti-tumor immunity, achieved through inhibition of processes such as MHC expression^11,71^, antigen processing and presentation and inflammatory cytokine production^72,73^. Moreover, our observation that NKG2D ligands are enriched in mesenchymal neuroblastomas coupled with the demonstration of the critical effector role of NK cells in the antitumor immune response against this tumor ^51^ and the promising responses of patients with other solid tumors to NKG2D-directed CAR NK cell therapy ^74^ strengthen this premise. Considered together, the results of our analysis identify cell lineage as an important determinant of the immune responsiveness of neuroblastoma and suggest rationales for the use of immune-based therapies, either alone or in combination with epigenetic inhibitors, against the two divergent phenotypes that define the lineage state of this pediatric tumor.

## Acknowledgements

We thank Matthew Harlow from the George lab, Sumit Sen Santara, Ying Zhang, and Zhibin Zhang from the Lieberman lab for helpful discussions. We thank Mark Zimmerman, A. Thomas Look and Kimberly Stegmaier for sharing cell lines. We are thankful to the following members of the former Haining lab at DFCI for sharing resources and experimental advice: Ulrike Gerdemann, Dawn Comstock, Kathleen Yates, Anna Word, and Adrienne Long. The results shown here are in part based on data curated by the R2: Genomics Analysis and Visualization Platform: http://r2.amc.nl/. This work was supported by a St. Baldrick’s Foundation Childhood Cancer Research Grant (R.E.G. and R.J.) DOD CA191000 (R.E.G. and R.J.) and NIH grants R01-CA197336; R01-CA148688 (R.E.G). St. S. is a recipient of the Pew Stewart Scholarship. Sa. S. and M.K. were supported by the Rally Foundation for Childhood Cancer Research and Infinite Love for Kids Fighting Cancer, B.C.M. by the National Center for Advancing Translational Sciences/NIH Award KL2 TR002542 and D.N.D. by an Alex’s Lemonade Stand Foundation Young Investigator Fellowship. This manuscript is dedicated to the memory of John R. Gilbert, Scientific Editor.

## Author contributions

Sa.S. and R.E.G. conceived the study. Sa.S., A.C., B.C.M., J.L., and R.E.G. designed the experiments. Sa.S. performed the molecular, cellular and genomic studies. S.D. conceived and performed the genomic and computational analysis with inputs from R.D., Sa.S. and R.E.G. Sa.S., A.C., and B.C.M. performed the T and NK cell studies. B.S. performed the animal and cloning experiments. S.Z. performed quantitative analysis of IHC images. H.H. performed the cell migration assays. M.K., D.N.D., and L.S. contributed to FACS analysis, generation of LDK378-resistant SH-SY5Y cells, and compound testing, respectively. M.C., R.V., R.J., and St.S. contributed ideas towards regulation of immune function and cell lineage state. Sa.S., S.D., and R.E.G. wrote the manuscript with input from all authors.

## Methods

### Cell culture

Human neuroblastoma (NB) cell lines (Kelly, NBL-S, CHP-212, SH-SY5Y, SH-EP, CHLA-20, NB69, SK-N-FI) were obtained from the Children’s Oncology Group cell line bank. ACN, GI-ME-N, NB-EbC1 were kind gifts from A. Thomas Look and Kimberly Stegmaier at Dana Farber Cancer Institute (DFCI). NB-9464 cells were provided by To-Ha Thai at Beth Israel Deaconess Medical Center, Boston, MA. The cell lines were authenticated through STR analyses at the DFCI Core facility and were routinely tested for mycoplasma. All NB cells were grown in RPMI-1640 medium (Invitrogen) supplemented with 10% fetal bovine serum (FBS) (Invitrogen) and 1% penicillin/streptomycin (Life Technologies). HEK293T cells obtained from the American Type Culture Collection (ATCC) were grown in DMEM (Invitrogen) supplemented with 10% FBS and 1% penicillin/streptomycin (Life Technologies). SH-SY5Y cells resistant to the ALK inhibitor ceritinib (LDK378) were described previously (Debruyne et al., 2016) and were grown in complete RPMI-1640 in the presence of 1.5 µM ceritinib.

### Generation of PRRX1-inducible cell lines

Lentiviral vectors containing wild type and DNA-binding mutants of PRRX1 were generated by cloning cDNAs encoding full length or homeodomain deletions of the human PRRX1A sequence into the pInducer20 lentiviral plasmid (gift from Stephen Elledge, Addgene plasmid #44012). The DNA-binding mutants harbor individual deletions of the three α-helices (ΔH1, ΔH2 and ΔH3) within the PRRX1 homeodomain (amino acids (aa) 94-153). Amino acid boundaries of the deleted regions are as follows: ΔH1 (aa 103-116); ΔH2 (aa 121-131); ΔH3 (aa 135-151). The lentivirus was packaged by co-transfection of pInducer20 plasmid with the helper plasmids, pCMV-deltaR8.91 and pMD2.G-VSV-G into HEK293T cells using the TransIT-LT1 Transfection Reagent (Mirus Bio LLC). Virus-containing supernatants were collected 48 hr after transfection. SH-SY5Y cells were transduced with the viral supernatant in the presence of 8 μg/ml polybrene (Sigma-Aldrich) and 24 hr later selected using neomycin (G418) (5 µg/ml). Induction of gene expression was achieved by treating cells every 2–3 days with doxycycline (dox; 200 ng/ml) in RPMI-1640 medium supplemented with 10% tetracycline-negative fetal bovine serum (tet-free FBS) (Gemini Bio-Products) and 1% penicillin/streptomycin.

### MYCN shRNA knockdown and IFN-y treatment

The pLKO.1 shRNA construct targeting MYCN (TRCN0000020694) was purchased from Sigma-Aldrich and the pLKO.1 GFP shRNA was a gift from D. Sabatini (Addgene plasmid #30323). Lentiviral packaging was performed in HEK293T cells as decribed above and viral supernatant was collected on days 2 and 3 after transfection. Kelly human NB cells were transduced with the viral supernatant in the presence of 8 μg/ml polybrene (Sigma-Aldrich) and 24 h later selected using puromycin (1 µg/ml) for 2 days. Puromycin-resistant Kelly cells were cultured for an additional 6 days. Cells were then treated with 100 ng/ml of recombinant human IFNγ (Biolegend) for 24h, following which they were harvested for analysis of RNA and protein expression.

### FACS analysis for cell surface protein staining

For each staining reaction 1 x 10^6^ live cells were placed in a 12 x 75 mm polystyrene round bottom tube (Falcon), resuspended in 100 µl 1x PBS and stained with the Zombie near-infrared (Zombie NIR) viability dye (BioLegend) at a 1:1,000 dilution for 15 minutes at RT. Cells were then washed once in FACS buffer (0.5% BSA in 1x PBS), resuspended in 100 µl of FACS buffer and incubated in 5 µl of Human TruStain FcX™ (Fc receptor blocking solution, BioLegend) for 10 minutes at RT. Next, appropriate volumes of conjugated fluorescent primary antibodies at predetermined optimum concentrations were added and incubated on ice for 20 minutes in the dark. Cells were then washed once in 2 ml of FACS buffer by centrifugation at 1500 rpm for 5 minutes. All FACS samples were analyzed on a FACSCalibur flow cytometer (Becton Dickinson) using Cell Quest software (Becton Dickinson). A minimum of 50,000 events was counted per sample and used for further analysis. Data were analyzed using FlowJo v10 software (Becton Dickinson). The following primary antibodies were used: PE-HLA (Biolegend; clone W6/32), PE-MICA/B (Biolegend clone 6D4), PE-ULBP2 (R&D Systems; clone 165903), PE-ULBP3 (R&D Systems; clone 166510), PE-H-2Kb (Biolegend AF6-88.5), PE-H-2Kb SIINFEKL (Biolegend; clone 25-D1.16), PE-mouse IgG2a k isotype control (Biolegend MOPC-173), PE-mouse IgG1 k isotype control (MOPC-21).

### Cell viability assay

SH-SY5Y parental and SH-SY5Y LDK-resistant cells were seeded in 96-well plates at a density of 2 × 10^3^ cells/well. After 24 h, cells were treated with increasing concentrations of LDK378 (ranging from 1 nM to 10 μM) dissolved in Dimethyl Sulfoxide (DMSO). DMSO solvent without the drug served as a negative control. Following 72 h incubation, cells were analyzed for viability using the CellTiter-Glo Luminescent Cell Viability Assay (Promega) according to the manufacturer’s instructions. Drug concentrations that inhibited cell growth by 50% (IC_50_) were determined using a non-linear regression curve fit with GraphPad Prism 8 software.

### Cell migration and invasion assays

Cell migration was measured using transwell chambers (Falcon). NB9464-H-2Kb^lo^ or NB9464-H-2Kb^hi^ cells in serum-free medium (0.5 x 10^6^ cells/ml) were added to the upper chamber and inserts (8 μm pore size) were placed in the lower chamber containing medium with 10% FBS. Following incubation at 37° C for 8 h, cells that migrated to the lower chamber were fixed with methanol and stained with crystal violet (Sigma-Aldrich). The stained cells were photographed with a light microscope at 100X magnification and migration was quantified as the number of cells per high power field. Cell invasion was measured using the fluorometric QCM^TM^ ECMatrix^TM^ Cell Invasion Assay (Millipore). NB9464-H-2Kb^lo^ or NB9464-H-2Kb^hi^ cells in serum-free medium (0.5 x 10^6^ cells/ml) were added to the upper chamber and inserts (8 μm pore size) placed in the lower chamber containing medium with 10% FBS. Following incubation at 37° C for 24 h, cell invasion was measured according to manufacturer’s instructions.

### EED226 treatment

5 x10^5^ SH-SY5Y cells were seeded into 10 cm plates and treated with either 5 µM EED226 (Selleck Chemicals) or DMSO (vehicle control) for 6-8 days, following which samples were harvested for downstream analyses. Cells were replenished with fresh media containing DMSO or EED226 every 2 days.

### Compounds

Ceritinib (LDK378) and EED226 were purchased from Selleck Chemicals. Doxycycline and dimethyl Sulfoxide (DMSO) was purchased from Sigma-Aldrich.

### RNA extraction and q-PCR

Total RNA was isolated using the RNAeasy Mini kit (Qiagen). Purified RNA was reverse transcribed to cDNA using Superscript IV VILO master mix (Thermo Fisher Scientific) following the manufacturer’s protocol. Quantitative PCR was performed using 1 µl cDNA, 1x PowerTrack SYBR Green PCR master mix (Thermo Fisher Scientific) and PCR primers (200 nM) in a total volume of 25 µl and analyzed on a ViiA 7 Real-Time PCR system (Thermo Fisher Scientific). Each individual biological sample was amplified in technical duplicate and normalized to GAPDH as an internal control. Relative expression was calculated according to the 2^−ΔΔCT^ quantification method (Livak and Schmittgen, 2001). PCR primer sequences are shown in Table S5.

### Synthetic RNA spike-in and RNA-sequencing

RNA-sequencing was performed on the following human NB cell lines: Kelly, NBL-S, CHP-212, SH-SY5Y, SH-SY5Y LDK-resistant and SH-EP. Biological duplicates (5 x 10^6^ cells per replicate) were homogenized in 1 ml of TRIzol Reagent (Invitrogen) and purified using the mirVANA miRNA isolation kit (Ambion) following the manufacturer’s instructions. Total RNA was treated with DNA-free^TM^ DNase I (Ambion), spiked-in with ERCC RNA Spike-In Mix (Ambion) and analyzed on an Agilent 2100 Bioanalyzer (Agilent Technologies) for integrity. Sequencing libraries were prepared using LP-KAPA mRNA Hyper Prep and sequenced using Illumina HiSeq for 40 bases.

### Western blotting

Cells were homogenized in NP40 lysis buffer (Life Technologies) containing 1× cOmplete EDTA-free protease inhibitor cocktail and 1x PhosSTOP (Roche). Protein concentration was measured using the DC Protein Assay (Bio-Rad). 100 µg total protein was denatured in LDS sample buffer (Invitrogen), separated on pre-cast 4-12% Bis-Tris gels (Invitrogen) and transferred to nitrocellulose membranes (Bio-Rad). Membranes were blocked using 5% dry milk (Sigma-Aldrich) in Tris-buffered saline (TBS) supplemented with 0.2% Tween-20 (TBS-T) for 1 hr, and incubated overnight with primary antibody in blocking buffer at 4 °C. Chemiluminescent detection was performed with appropriate HRP-conjugated secondary antibodies and enhanced chemiluminescence reagents (Thermo Scientific). Images were developed using Genemate Blue ultra-autoradiography film (VWR).

### Antibodies

The following primary antibodies were used: MYCN (Cat #51705), MYC (13987), GATA3 (5852), TAP1 (12341), TAP2 (12259), LMP7 (13635), NOTCH1 (3608), cleaved NOTCH1 (4147), SOX9 (82630), AXL (8661), GAPDH (2118), β-actin (3700), IRF1 (8478), VIM (5741), YAP1 (4912), TAZ1 (4883) (Cell Signalling Technologies (CST)); PHOX2B (Abcam;183741), FN1 (RnD systems; AF1918), LMP2 (Santa Cruz; 271354) and PRRX1 (Santa Cruz; 293386).

### Chromatin immunoprecipitation-quantitative PCR (ChIP-qPCR)

Soluble chromatin was prepared as above from SH-SY5Y cells without or with dox-inducible PRRX1 expression (200 ng/ml dox for 10 days). ChIP was performed as described in the preceding section using the following antibodies: H3K4me3 (Abcam 8580), H3K27me3 (Millipore 07-729), EZH2 (CST 5246), SUZ12 (CST 3737), EED (Millipore 17-10034), rabbit IgG (CST 2729). Purified ChIP DNA was dissolved in 60 µl of 1x TE. Quantitative PCR was performed on a ViiA 7 Real-Time PCR system (Thermo Fisher Scientific) with 1 µl purified DNA, 1x PowerTrack SYBR Green PCR master mix (Thermo Fisher Scientific) and PCR primers (200 nM) against the genomic regions of interest. Each individual biological sample was amplified in technical duplicate. Relative enrichment was quantified using the percent input method. PCR primer sequences are shown in Table S5.

### Chromatin immunoprecipitation-sequencing (ChIP-seq)

Approximately 10-12 x 10^7^ cells were crosslinked with 1% formaldehyde (Thermo Scientific) for 10 min at room temperature (RT) followed by quenching with 0.125 M glycine for 5 min. The cells were then washed twice in ice-cold 1x Phosphate Buffered Saline (PBS), and the cell pellet equivalent of 4 x 10^7^ cells were flash frozen and stored at −80°C. Crosslinked cells were lysed in lysis buffer 1 (50 mM HEPES-KOH pH7.5, 140 mM NaCl, 1 mM EDTA, 10% glycerol, 0.5% NP40, 0.25% Triton X-100). The resultant nuclear pellet was washed once in lysis buffer 2 (10 mM Tris-HCl pH 8, 200 mM NaCl, 1 mM EDTA, 0.5 mM EGTA) and then resuspended in sonication buffer (50 mM HEPES-KOH pH 7.5, 140 mM NaCl, 1mM EDTA, 1mM EGTA, 1% Triton X-100, 0.1% sodium deoxycholate, 0.2% SDS). Chromatin was sheared using a Misonix 3000 sonicator (Misonix) and at the following settings: 10 cycles, each for 30 s on, followed by 1 min off, at a power of approximately 20 W. The lysates were then centrifuged for 15 min at 4 °C, supernatants collected and diluted with an equal amount of sonication buffer without SDS to reach a final concentration of 0.1% SDS. For each ChIP, the chromatin equivalent of 1 x 10^7^ cells was used. 50 µl of Protein G Dynabeads per sample (Invitrogen) were blocked with 0.5% BSA (w/v) in 1x PBS. Magnetic beads were loaded with the following antibodies: 10 µg of H3K27me3 (Millipore 07-729); 3 µg of H3K27ac (Abcam 4729), and 3 µg of H3K4me3 (Abcam 8580) and incubated overnight at 4°C. The sonicated lysates were then incubated overnight at 4°C with the antibody-bound magnetic beads, washed with low-salt buffer (50 mM HEPES-KOH (pH 7.5), 0.1% SDS, 1% Triton X-100, 0.1% sodium deoxycholate, 1 mM EGTA, 1 mM EDTA, 140 mM NaCl and 1× complete protease inhibitor), high-salt buffer (50 mM HEPES-KOH (pH 7.5), 0.1% SDS, 1% Triton X-100, 0.1% sodium deoxycholate, 1 mM EGTA, 1 mM EDTA, 500 mM NaCl and 1× complete protease inhibitor), LiCl buffer (20 mM Tris-HCl (pH 8), 0.5% NP-40, 0.5% sodium deoxycholate, 1 mM EDTA, 250 mM LiCl and 1× complete protease inhibitor) and Tris-EDTA (TE) buffer. DNA was then eluted in elution buffer (50 mM Tris-HCl (pH 8.0), 10 mM EDTA, 1% SDS), and high-speed centrifugation performed to pellet the magnetic beads and collect the supernatants. The crosslinking was reversed overnight at 65° C in the presence of 300 mM NaCl. RNA and protein were digested using RNase A and proteinase K, respectively, and DNA was purified with phenol-chloroform extraction and ethanol precipitation. Purified ChIP DNA was used to prepare Illumina multiplexed sequencing libraries using the NEBNext Ultra II DNA Library Prep kit and the NEBNext Multiplex Oligos for Illumina (New England Biolabs) according to the manufacturer’s protocol. Libraries with distinct indices were multiplexed and run together on the Illumina NextSeq 500 (SY-415-1001, Illumina) for 75 base pairs.

### IFN-gamma induction and antigen presentation in NB9464 cells

Approximately 1 x 10^6^ cells were seeded onto 10 cm plates. 24 hr later, adherent cells were treated with recombinant mouse IFN-γ (Biolegend) for 24h and harvested for H-2Kb analysis using FACS as described above. For antigen presentation assays, cells treated with IFN-γ were pulsed with SIINFEKL (OVA peptide) at 37°C. Cells were subsequently washed with 1x PBS to remove unbound peptide and processed for analysis using FACS.

### T cell activation assays

OT-I T cell receptor (TCR) transgenic mice were purchased from Jackson Laboratories (Bar Harbor, ME). Splenocytes were harvested and T cells were subsequently isolated from the mononuclear layer using Ficoll separation and directly used in co-culture assays. Successful enrichment of CD8^+^ T cells was confirmed by FACS analysis using the FITC-CD8 antibody. Cells were pulsed with SIINFEKL (OVA peptide) at 37 °C, to bind to cell surface H-2Kb. Cells were subsequently washed with 1x PBS to remove unbound peptide and then co-cultured with unstimulated OT-1 T cells for 24 h. OT-I cells were then harvested, and sequentially stained with the Zombie NIR viability dye and FITC-CD8, PE-CD69 antibodies, followed by fixation with 1% paraformaldehyde (Polysciences, Inc). OT-1 cells were analyzed by flow cytometry using a FACSCanto II cell analyzer (Becton Dickinson) and FlowJo V10 software (Becton Dickinson).

### In vitro assays for NKG2D binding

SH-SY5Y parental, SH-SY5Y LDK-resistant, and SH-EP cells were incubated with recombinant human NKG2D-Fc chimeric protein or an equivalent concentration of human IgG, following which cells were washed, and sequentially stained with the Zombie NIR viability dye as described above followed by incubation with an Alexa 647-conjugated anti-human IgG antibody for 30 minutes. Cells were washed in FACS buffer and analyzed by flow cytometry using a FACSCanto II cell analyzer (Becton Dickinson) and FlowJo V10 software (Becton Dickinson).

### NK cell degranulation assays

Human peripheral blood NK cells were isolated from blood collars using a RosetteSep™ human NK cell enrichment cocktail (STEMCELL Technologies). NK cells were then co-cultured for 4 h with confluent monolayers of SH-SY5Y parental, SH-SY5Y LDK-resistant, and SH-EP cells at in the presence of CD107a antibody (Biolegend). Additionally, co-cultures of NK cells and 721.221 B cells were included as positive controls for degranulation. At the endpoint, NK cells were harvested, stained with Zombie Yellow (Biolegend) and CD56 FITC (or NKp46 AlexaFluor 647^TM^) and NKG2D PE (or mouse IgG1 PE) antibodies (Biolegend), followed by fixation with 1% paraformaldehyde (Polysciences, Inc). For the NKG2D blocking assay, NK cells were incubated with purified anti-NKG2D antibody or mouse IgG1 isotype control at 37 °C, following which the degranulation assay was performed as detailed above. NK cells were analyzed by flow cytometry using a FACS Canto II (Becton Dickinson) and FlowJo V10 software (Becton Dickinson).

### Data sets

Publicly available RNA-seq data (GEO accession number GSE49711/GSE62564) from a cohort of 498 primary human neuroblastoma tumors, microarray expression data from 394 neuroblastoma tumors (GSE120572) and 24 human neuroblastoma cell lines (GSE28019) were accessed through the R2 genomics analysis and visualization platform (https://hgserver1.amc.nl/cgi-bin/r2/). Clinical annotations for tumors were obtained from GSE49711/GSE62564 regarding MYCN status (MYCN-nonamplified vs. MYCN-amplified, INSS stage [high (stage 4) vs. low (1, 2, 3 and 4s], risk status (high vs. low) and age (< 18 months vs. ≥ 18 months).

### Analysis of RNA-sequencing data

#### RNA-seq data processing and identification of differentially expressed genes

Single-end RNA-seq samples with 40 base pair (bp) read lengths were mapped to the human genome (GRCh38) and ERCC spike-in sequences. Reads were mapped to the genome using Bowtie2 (version 2.3.4.3) and default parameters. Reads that overlapped with the genomic location for exonic regions were used to calculate gene counts with the FeatureCounts (Subread package of version 1.6.3) package. Further, spike-in read counts were used for each sample to normalize the library sizes. These read counts were used to calculate the sample-specific size factor by using the function estimateSizeFactors (DESeq2) available in R. Normalized sample coverage profiles were then created from previously determined size factors by using bamCoverage (DeepTools v3.0.2) and parameters “--scaleFactor --skipNonCoveredRegions”. To check the reproducibility of biological replicates for each condition, principal component analysis (PCA) and correlation (Spearman’s rank coefficient) were assessed from the sample coverage profiles at genome-wide scale and visualized using scatterplots and heatmaps. Because these analyses showed a high correlation of sample coverage profiles between replicates, replicates were merged using samtools merge and processed again as described for the individual replicates. Next, differential gene expression analysis was performed using the DESeq2 in R. To detect differentially expressed genes (DEGs) in each sample, raw read counts from RNA-seq data were imported to the DESeq2 and the size factors calculated using the estimateSizeFactors function. A transcript with an absolute log2 fold-change ≥ 1.5 and an adjusted *P*-value ≤ 0.01 was considered significant.

#### Enrichment analysis

Gene ontology enrichment for selected gene sets was performed by the Enrichr program (https://amp.pharm.mssm.edu/Enrichr/). All GO terms were ranked based on the Enrichr combined score, calculated by multiplying the adjusted *P*-value with the z-score using the Fisher’s exact test. The Fisher’s exact test was used to determine significant overlaps between the queried gene sets and other publicly available datasets. Enrichment of gene sets was considered significant for an adjusted P-value ≤ 0.01, unless stated otherwise.

#### Estimation of immune cell content in neuroblastoma tumors

Cell type identification by estimating relative subsets (CIBERSORT) ^43^, a deconvolution method was used to evaluate immune cell fractions from gene expression data using the R package ‘immunedeconv’. RNA transcript estimations were generated for all 498 neuroblastoma tumors using the LM22 signature matrix available for 22 immune cell types. CIBERSORT was run in “Absolute mode” with disabled quantile normalization as recommended for tumor RNA-seq data and the overall immune content produced by the algorithm compared among tumors.

### ChIP-seq analysis

#### Data processing

All ChIP-seq raw datasets were processed as previously described ^83^. The raw read quality of the samples was accessed using the Fastqc tool (v0.11.7) to identify possible sequencing errors and biases. Reads were aligned to the human genome (build hg19, GRCh37.75) using the mapper Bowtie (v2.3.4.3) with default parameters. Unique and non-duplicate reads that mapped to the reference genome were further processed using Samtools (v1.9) and the MarkDuplicates (v2.1.1) command of Picard tools. Next, antibody enrichment in each replicate as compared to input samples was verified using the PlotFingerprint command of deepTools (v3.1.1). Peak caller MACS2 (2.1.1) was used to identity narrow peaks (H3K4me3 and H3K27ac) with the parameters “--q 0.01--call-summits” and broad peaks (H3K27ac and H3K27me3) with the parameters “--broad-cutoff 0.01”. Peaks that overlapped with black-listed regions (http://mitra.stanford.edu/kundaje/akundaje/release/blacklists/) of the reference genome (mostly comprised of major satellite repeats of telomeric and pericentromeric regions) were filtered out. Command “bamCompare” from the deepTools was used with the parameters “--scaleFactorsMethod readCount --binSize 40 --operation subtract --smoothLength 80 -- extendReads 200” to create the input normalized bedgraph tracks for each replicate and afterwards negative values were set to zero and counts were scaled to reads per million/base pair (rpm/bp) to account for differences in the library size. Bigwig files were created for visualization with bedGraphToBigWig. Subsequently, correlations among the ChIP-seq replicates were accessed using bigwigs with the command “multiBigwigSummary” from deepTools and highly correlated replicates merged at the BAM level. Peak identifications were then repeated in the same manner for these merged BAM files.

#### Identification of super-enhancer regions

Super-enhancers (SEs) were identified using the ROSE algorithm (https://bitbucket.org/young_computation/rose/src/master/). Briefly, H3K27ac binding regions identified by MACS2 as significant peaks, termed typical enhancers, were stitched together if they were within 12.5 kb of each other. These stitched enhancers were ranked by comparing the H3K27ac signal (density * length) with the input signal. The ROSE algorithm was used to determine the inclination point for all stitched H3K27ac signals and to segregate regular enhancers from SEs. To compare SEs in 5Y-parental, 5Y-LDK-resistant and SH-EP cells, the same maximum threshold was used between the conditions.

#### Analysis of histone binding changes between lineage states

To analyze the changes in occupancies of active (H3K27ac and H3K4me3) and repressive (H3K27me3) histone marks during the transition from adrenergic (5Y-par.) to mesenchymal (5Y-LDK-res., SHEP) states, we compared the peaks of histone marks identified by MACS2 at the promoter regions. For this purpose, we first extracted the promoter regions ± 2 kb with respect to the TSS (−2kb upstream to +2kb downstream) of all annotated protein coding genes and subsequently, retrieved the peaks of H3K27ac, H3K4me3 and H3K27me3 from 5Y-par., 5Y-LDK-res. and SH-EP cells. Now, to determine the differential binding of each histone mark between 5Y-par. and 5Y-LDK-res. cells, we first combined all significant peaks called by MACS2 at the promoter regions and merged the peak regions that overlapped by at least 50%. This 50% threshold was used to avoid merging peaks that had clear and distinct summits. Next, the normalized active or repressive histone marks read densities were calculated for each region and a ratio of [log2 (5Y-LDK-res./ 5Y-par.)] was calculated. Shared peaks had similar enrichment of either active or repressive histone marks in both the cell types. Similar comparisons were made for active or repressive histone marks between 5Y-par. and SH-EP cells. To further compare changes in all histone marks at the promoters of immune genes, the gain of significant H3K27ac, H3K4me3 binding and loss of H3K27me3 signals were listed in mesenchymal (5Y-LDK-res., SHEP) cells as compared to adrenergic (5Y-par.) cells.

#### Integrated analysis of histone binding and gene expression

Cell-type specific differential enrichment of H3K27me3 and H3K4me3 binding in 5Y-par., 5Y-LDK-res. and SH-EP cells was determined by calculating the log2 (H3K4me3+1/H3K27me3+1) ratios in the promoter regions (TSS ± 2 kb) of immune genes. Next, to examine the association between gene expression and differential enrichment of H3K27me3 and H3K4me3 binding in immune genes in 5Y-par., 5Y-LDK-res. and SH-EP cells, genes were ranked based on their expression values and plotted against the calculated ratios between the histone marks.

---

**Supplementary Fig. 1.**
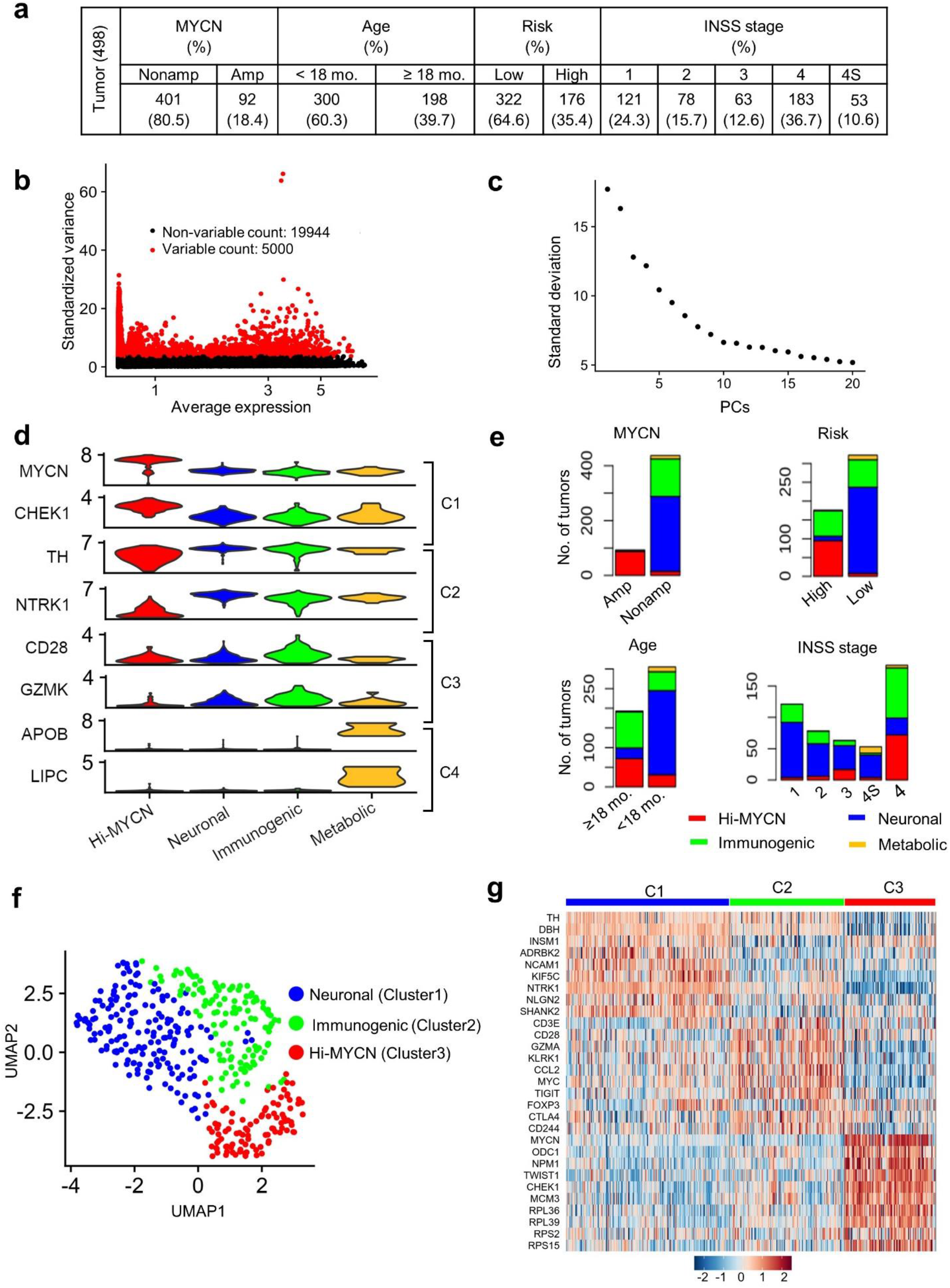
Tumors within the immunogenic cluster express both immune activation and evasion markers. **(a)** Distribution of the 498 primary NB tumors in the data set (SEQC-498; GSE49711) within the indicated prognostic categories. **(b)** Scatter plot of the standardized variance in expression of all protein coding genes within 498 tumors. Red dots indicate the top 5000 variably expressed genes. **(c)** Elbow plot representing the percentage variance for the top 20 principal components, PCs. **(d)** Violin plots showing the expression of representative marker genes across the four clusters. **(e)** Stacked bar plots showing the distribution of tumors within the defined prognostic features within each cluster. Amp, amplified; Nonamp, nonamplified. **(f)** Two-dimensional UMAP representations of the gene expression profiles in 394 NB tumors (GSE120572). Each dot represents a tumor. The top 3000 highly variable genes were selected based on the variance-stabilizing method ^34^ and the 20 significant principal components (PCs) selected and processed in UMAP to generate three clusters representing three NB subtypes. The DEGs were identified for each cluster using the receiver operating characteristics (ROC) curve to compare one cluster with other two (log_2_ FC > 0.25). **(g)** Heat map of expression values of 10 representative DEGs within each cluster. Rows are z-score scaled average expression levels for each gene in all three clusters.

**Supplementary Fig. 2.**
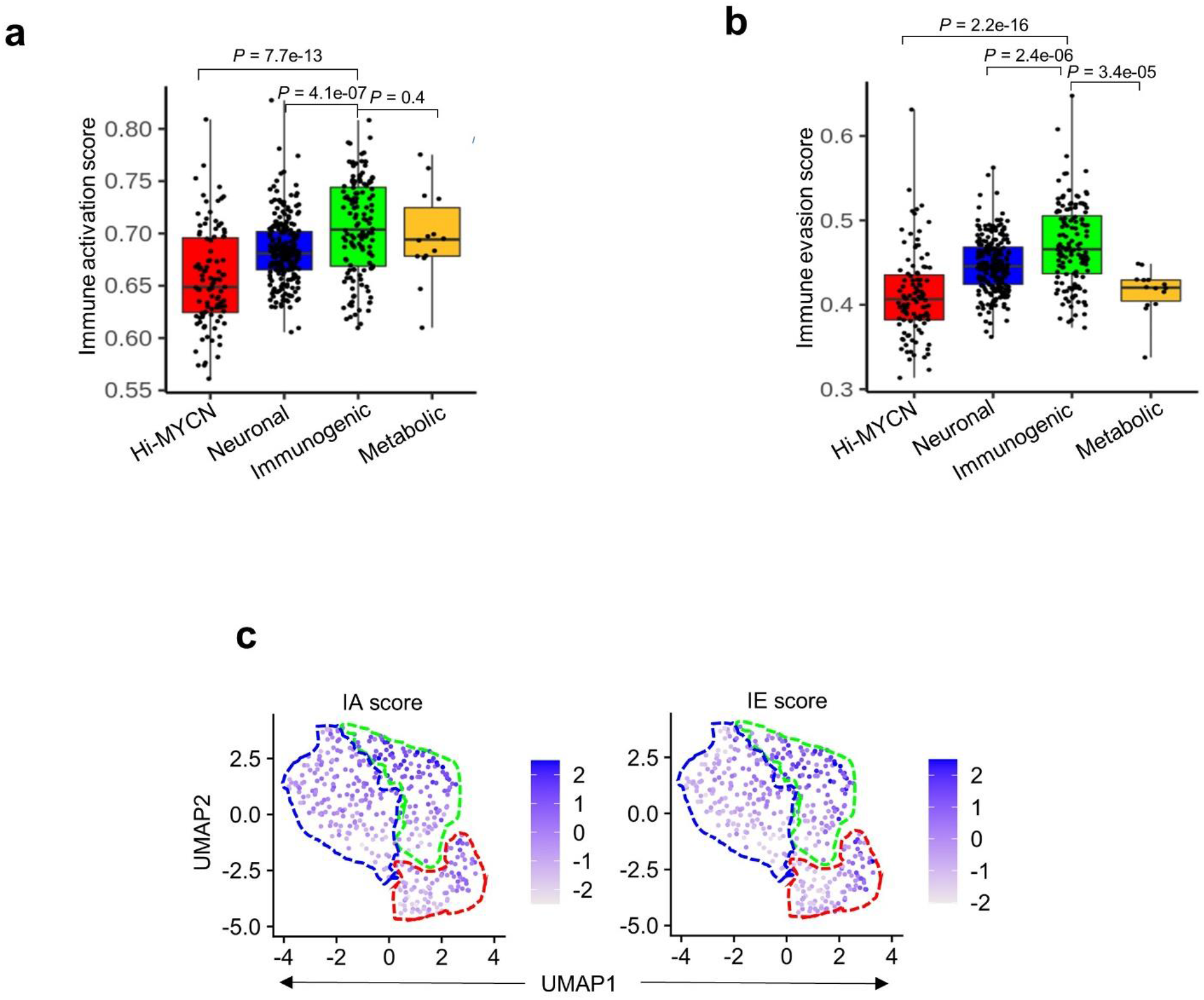
The immunogenic tumors are associated with markers of both immune activation and evasion. **(a, b)** Box plots comparing immune activation **(a)** and evasion **(b)** scores within the four clusters from the SEQC-498 tumor data set. All box plots are defined by center lines (medians), box limits (25^th^ and 75^th^ percentiles), whiskers (minima and maxima; the smallest and largest data range). Significance was determined by the Wilcoxon rank-sum test. **(c)** UMAP visualization of the distribution of IA and IE scores among the three tumor clusters derived from the 394 NBs in the GSE120572 dataset. Color bar represents normalized z-scores. Values <2.5 and >2.5 were set to −2.5 and +2.5 respectively, to reduce the effects of extreme outliers.

**Supplementary Fig. 3.**
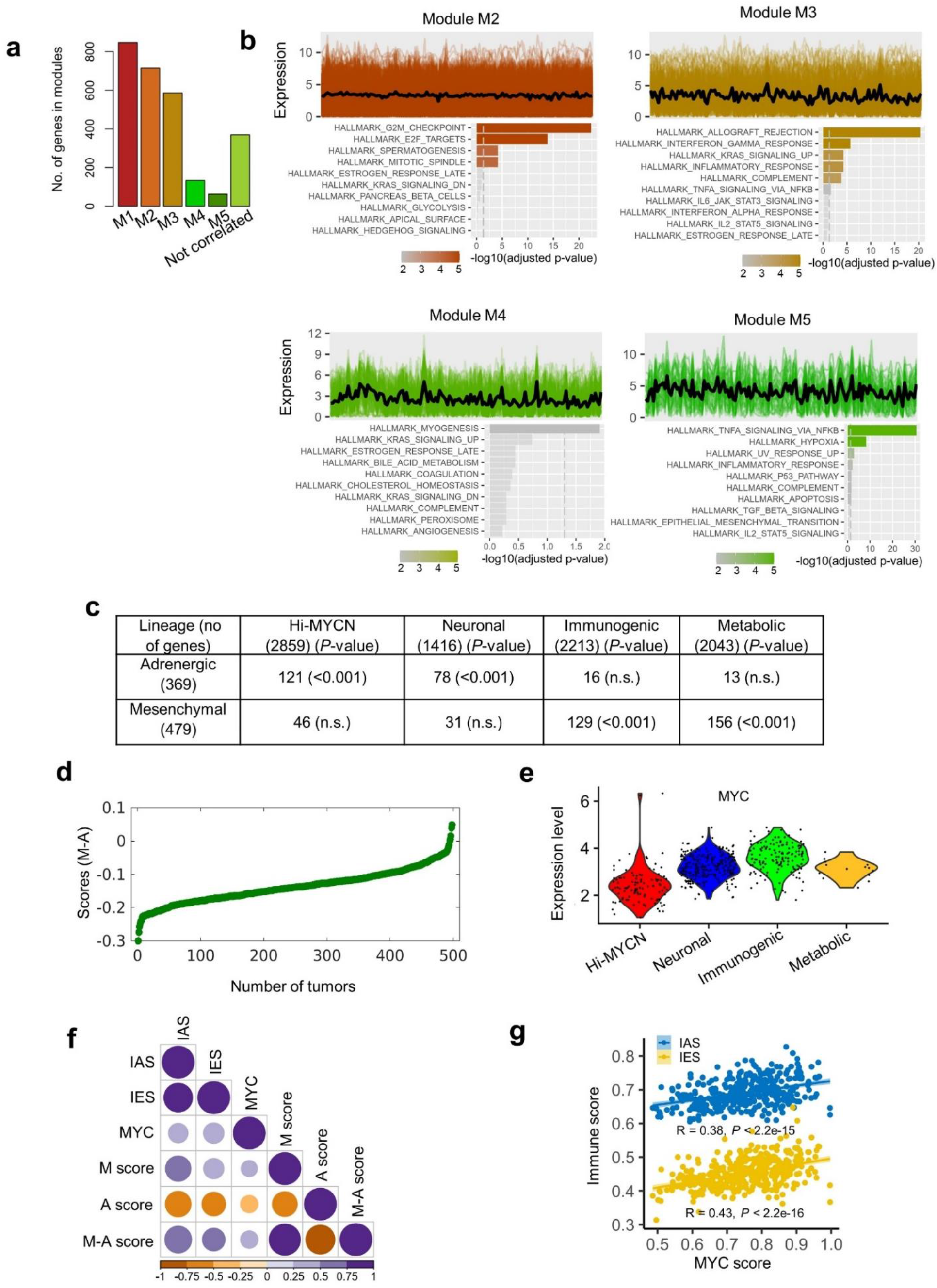
Cell lineage markers are significantly associated with immune gene signatures in NB. **(a)** Bar plots representing the numbers of co-expressed genes within each module. **(b)** Profile plots depicting the expression levels (y-axis) of individual genes (colored lines) and their mean expression (black line) in 140 tumors (x-axis) from modules M2, M3, M4 and M5. GO analysis of co-expressed genes associated with each module using the KEGG database is appended below the respective profile plot. The vertical dashed line indicates the adjusted *P*-value of 0.05. **(c)** Summary of the overlap between the DEGs associated with the four tumor clusters and the adrenergic or mesenchymal signature genes as per Groningen et al., 2017. Significance was determined by Fisher’s exact test. **(d)** Scatter plot of the 498 primary NB tumors ranked based on increasing M-A score. **(e)** Violin plots of the distribution of normalized expression levels of MYC in the four tumor clusters. **(f)** Pearson correlation matrix showing pairwise correlation values among the indicated parameters. The colors and sizes of the circles indicate the correlation coefficient values, with the least (smaller, orange circles) to the most (larger, blue circles) degree of association between the parameters shown. **(g)** Scatter plot of the correlation between MYC expression and immune activation (blue dots) or evasion (yellow dots) scores in *MYCN-*nonamplified tumors.

**Supplementary Fig. 4.**
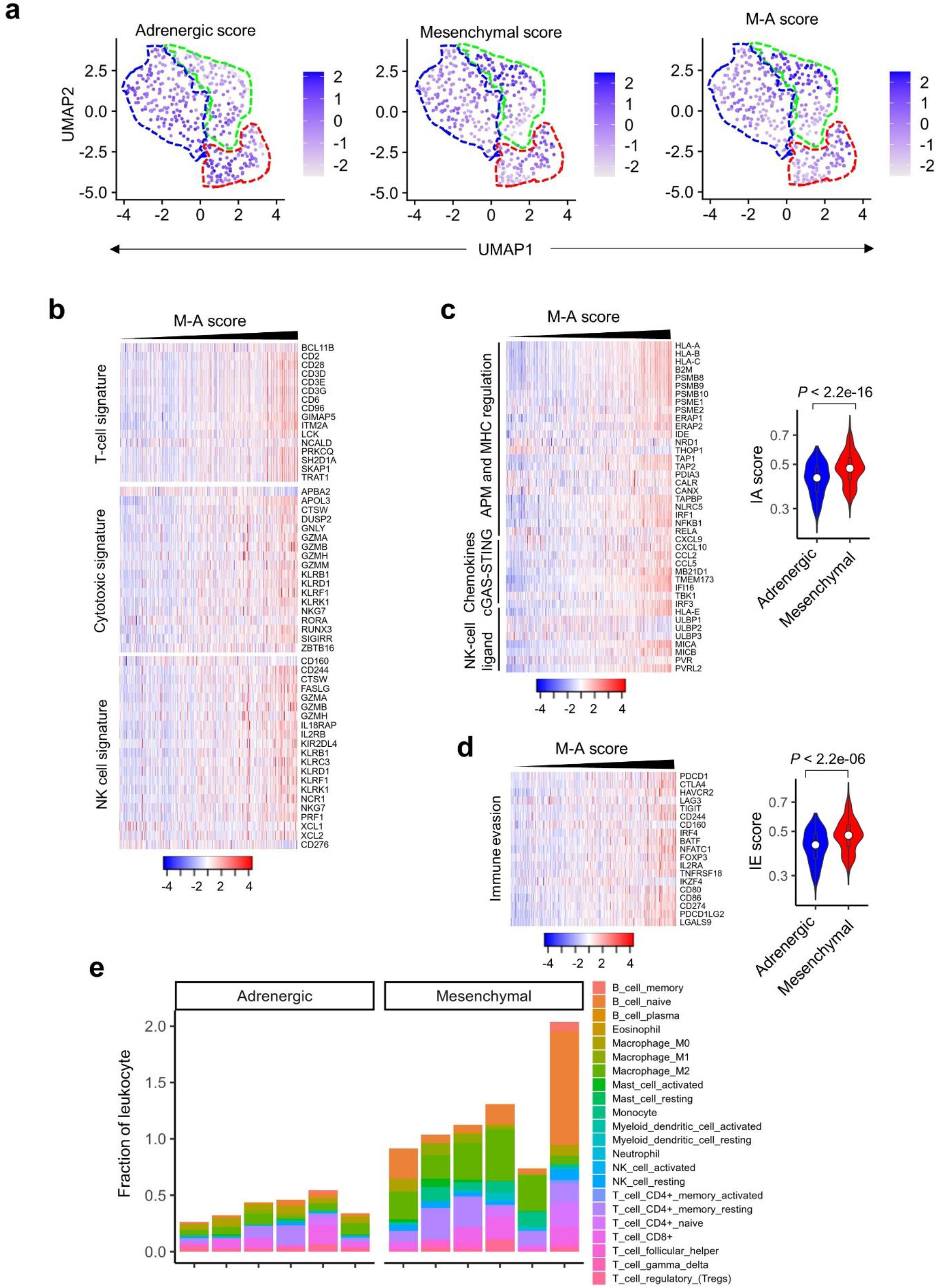
The relative mesenchymal score (M-A score) is positively correlated with an immunogenic signature. **(a)** UMAP visualization of the distribution of adrenergic, mesenchymal, and M-A scores among the three tumor clusters derived from 394 NBs in the GSE120572 dataset. Color bar represents normalized z-scores. Values <2.5 and >2.5 were set to −2.5 and +2.5 respectively, to reduce the effects of extreme outliers. **(b)** Heat maps of the indicated immune cell signatures in *MYCN-*nonamplified tumors, ranked by increasing M-A scores. Log_2_ gene expression values were z-score transformed for heatmap visualization. **(c, d)** Heatmaps depicting the immune activation **(c)** and evasion **(d)** signatures in *MYCN-*nonamplified tumors, ranked by increasing M-A score. Log_2_ gene expression values were z-score transformed for visualization. Violin plots comparing the distribution of immune activation **(c)** and evasion **(d)** signatures in 100 tumors from upper (mesenchymal) and lower (adrenergic) quartiles of the M-A score are shown next to the heatmaps. Significance determined by the two-sided KS test. Box plots within the violin plots are defined by center lines (medians), box limits (25^th^ and 75^th^ percentiles), whiskers (minima and maxima; 1.5X the interquartile range). **(e)** Bar graph comparing the CIBERSORT-estimated fractional content of the indicated tumor-infiltrating leukocytes in six tumors with the uppermost (mesenchymal) and lowermost (adrenergic) M-A scores.

**Supplementary Fig. 5.**
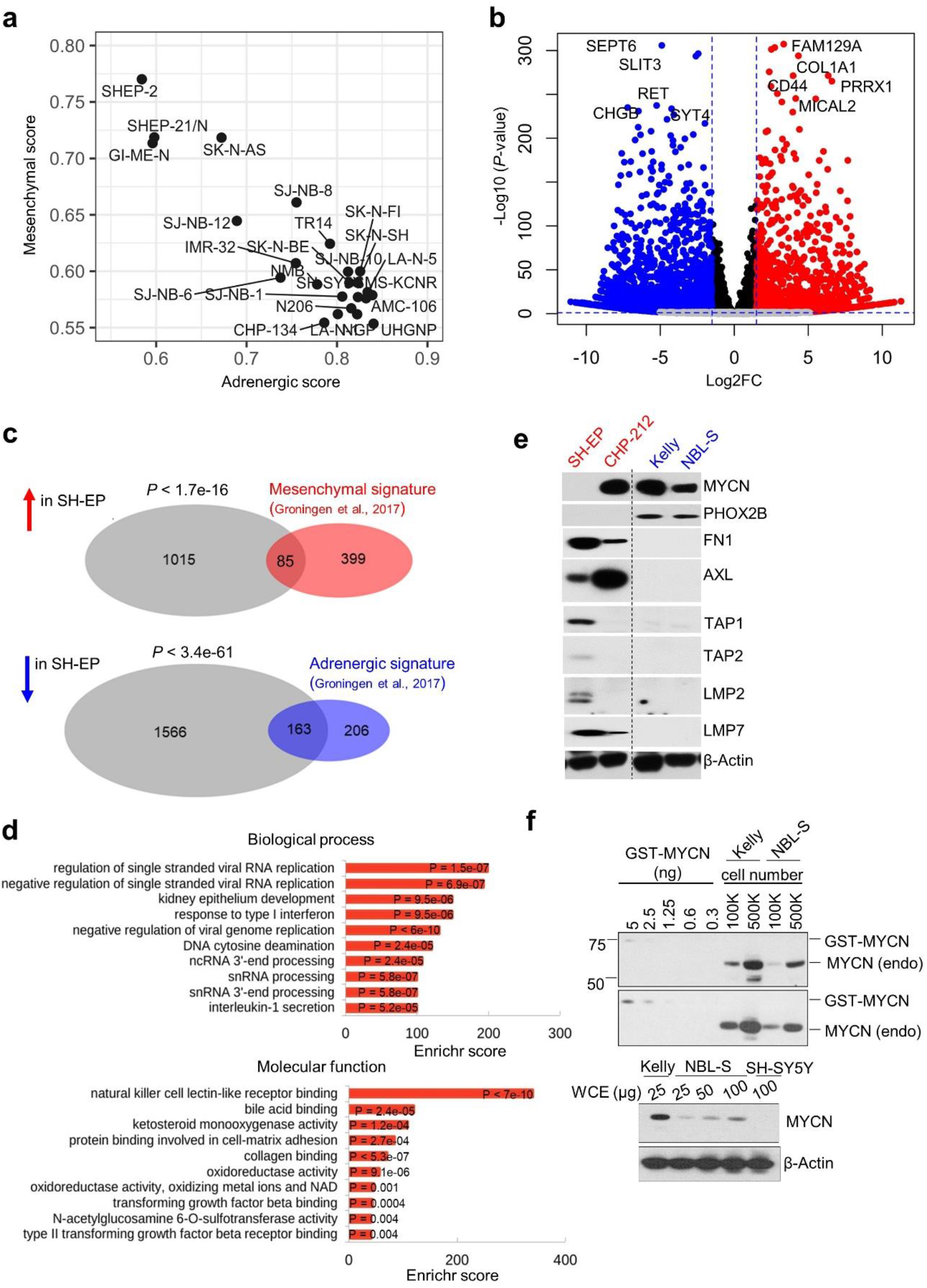
Mesenchymal lineage-specific marker and cell-intrinsic immune gene expression are significantly correlated in NB. **(a)** Scatter plot of adrenergic and mesenchymal scores in NB cell lines (GSE28019). **(b)** Volcano plot showing the gene expression changes between mesenchymal SH-EP and adrenergic SH-SY5Y cells. The top ten lineage marker genes are highlighted. The fold changes are represented in log_2_ scale (X-axis) and the - log_10_ of the *P-*values depicted on Y-axis (FDR < 0.1 and log_2_FC > 1). **(c)** Venn diagram of the overlap between DEGs in SH-EP cells (compared to SH-SY5Y cells) and the mesenchymal or adrenergic signatures derived from Groningen et al^38^. Statistical significance was determined using Fisher’s exact test. **(d)** GO analysis of differentially upregulated genes in mesenchymal SH-EP compared to adrenergic SH-SY5Y cells. **(e)** WB analysis of cell lineage marker and antigen processing gene expression in the indicated NB cells. Dotted line indicates the margin where gel images have been cut. **(f)** *Upper*, WB analysis of MYCN levels in the indicated numbers of Kelly and NBL-S NB cells titrated against known amounts of purified GST-MYCN protein. *Lower*, WB analysis comparing MYCN levels in whole cell extracts (WCE) from Kelly cells to titrated levels from WCE in NBL-S cells. SH-SY5Y cells that do not express MYCN serve as a negative control. Actin was used as a loading control in **(e)** and **(f)**.

**Supplementary Fig. 6.**
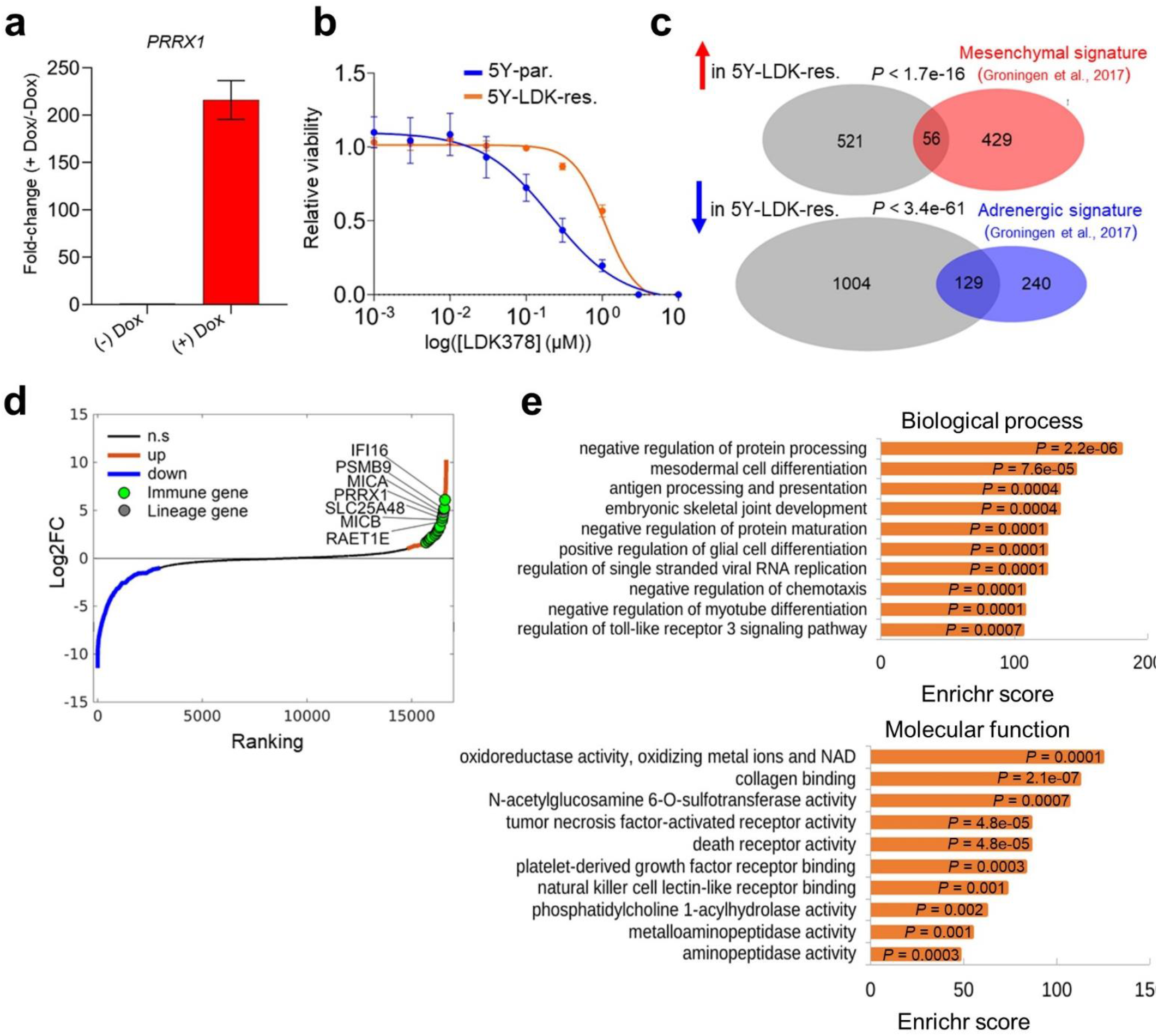
Acquired resistance to ceritinib (LDK387) in adrenergic SH-SY5Y cells is associated with reprogramming to the mesenchymal lineage and increased expression of immune response genes. **(a)** RT-qPCR analysis of PRRX1 expression in adrenergic SH-SY5Y cells engineered to express doxycycline (dox)-inducible PRRX1 in the presence or absence of dox (200 ng/mL) for 10 days. Data represent the means ± SD, *n* = 2 biological replicates. **(b)** Dose–response curves of ceritinib (LDK378)-sensitive (5Y-par.) and - resistant (5Y-LDK-res.) SH-SY5Y cells treated with increasing concentrations of LDK378 for 72 h. Data represent means ± SD, *n* = 2 biological replicates. **(c)** Venn diagrams depicting the overlap between the DEGs in LDK-resistant SH-SY5Y cells (compared to parental SH-SY5Y cells) and the mesenchymal or adrenergic signatures derived from Groningen et al^38^. *P*-values were determined by Fisher’s exact test. **(d)** Waterfall plot of the fold-change in RNA expression levels of up- and down-regulated genes in LDK-resistant SH-SY5Y cells compared to parental SH-SY5Y cells; selected immune genes are highlighted in green. **(e)** GO analysis of differentially upregulated genes in LDK-resistant SH-SY5Y compared to parental SH-SY5Y cells.

**Supplementary Fig. 7.**
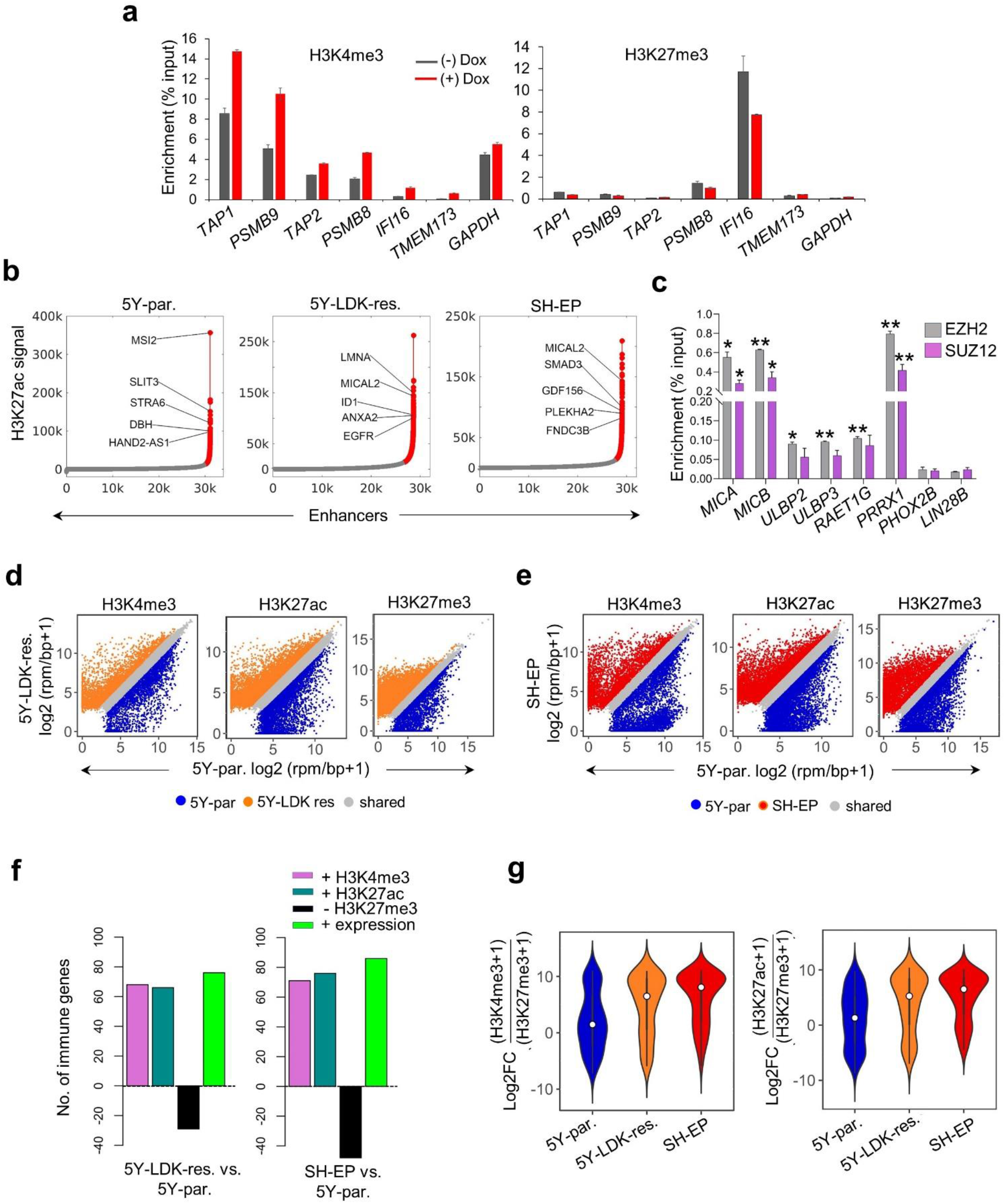
Immune gene activation associated with the mesenchymal cell state is epigenetically regulated. **(a)** ChIP-qPCR analysis of H3K4me3 and H3K27me3 enrichment at the promoters of the indicated immune genes in SH-SY5Y cells expressing doxycycline-inducible PRRX1 in the presence or absence of dox (200 ng/mL) for 10 days. Enrichments at TAP1 and PSMB9 loci correspond to amplicons 4 and 6 respectively, as described in Fig. 5a. Data represent the means ± SD, *n* = 2 biological replicates. **(b)** Identification of enhancer regions in parental SH-SY5Y, LDK-resistant SH-SY5Y and SH-EP cells. H3K27ac bound regions identified as significant peaks were stitched together if they were within 12.5 kb of each other and termed typical enhancers (plotted in grey). Super enhancers (SEs) were defined as stitched enhancers surpassing the threshold signal based on the inclination point in all cell types (plotted in red). In parental SH-SY5Y, LDK-resistant and SH-EP cells, 2.94% (915/31116), 6.56% (1880/28635) and 4.18% (1215/29057) of the enhancers were classified as SEs respectively. The top five SE-associated lineage-specific genes are highlighted. **(c)** ChIP-qPCR analysis of EZH2 and SUZ12 enrichment at the indicated genes in adrenergic 5Y-par. cells. Data represent the means ± SD, *n* = 2 biological replicates, \**P* < 0.05; \*\**P* < 0.01 two-tailed Student’s t-test. *P*-values were calculated in comparison to enrichment observed at the Lin28B TSS (negative control locus). **(d, e)** Scatter plots representing the differential binding of the indicated histone marks at the promoter regions (TSS ± 2 kb) of all protein coding genes between parental SH-SY5Y (5Y-par.) and LDK-resistant SH-SY5Y (5Y-LDK res.) **(d)**, and parental SH-SY5Y and SH-EP cells **(e)**. rpm/bp, reads per million mapped reads per base pair. A ≥ 0.75 log_2_FC threshold was used to identify unique peaks for each individual histone mark. Unique and shared peaks are shown in different colors. **(f)** Bar plots representing the numbers of immune genes with increased deposition of H3K4me3 and H3K27ac (log_2_ FC ≥ 0.75, TSS ± 2 kb) and loss of H3K27me3 (log_2_ FC ≥ 0.75, TSS ± 2 kb) histone marks, together with increased RNA expression (log_2_ FC ≥ 1) in mesenchymal LDK-resistant SH-SY5Y (*left*) or SH-EP (*right*) as compared to adrenergic parental SH-SY5Y cells. **(g)** Violin plots of the ratios of active to repressive histone marks (*left*, H3K4me3:H3K27me3; *right*, H3K27ac:H3K27me3) surrounding immune gene promoters (TSS ± 2kb) in parental SH-SY5Y, LDK-resistant SH-SY5Y and SH-EP cells. Significance was determined by the two-sided Wilcoxon rank-sum test.

**Supplementary Fig. 8.**
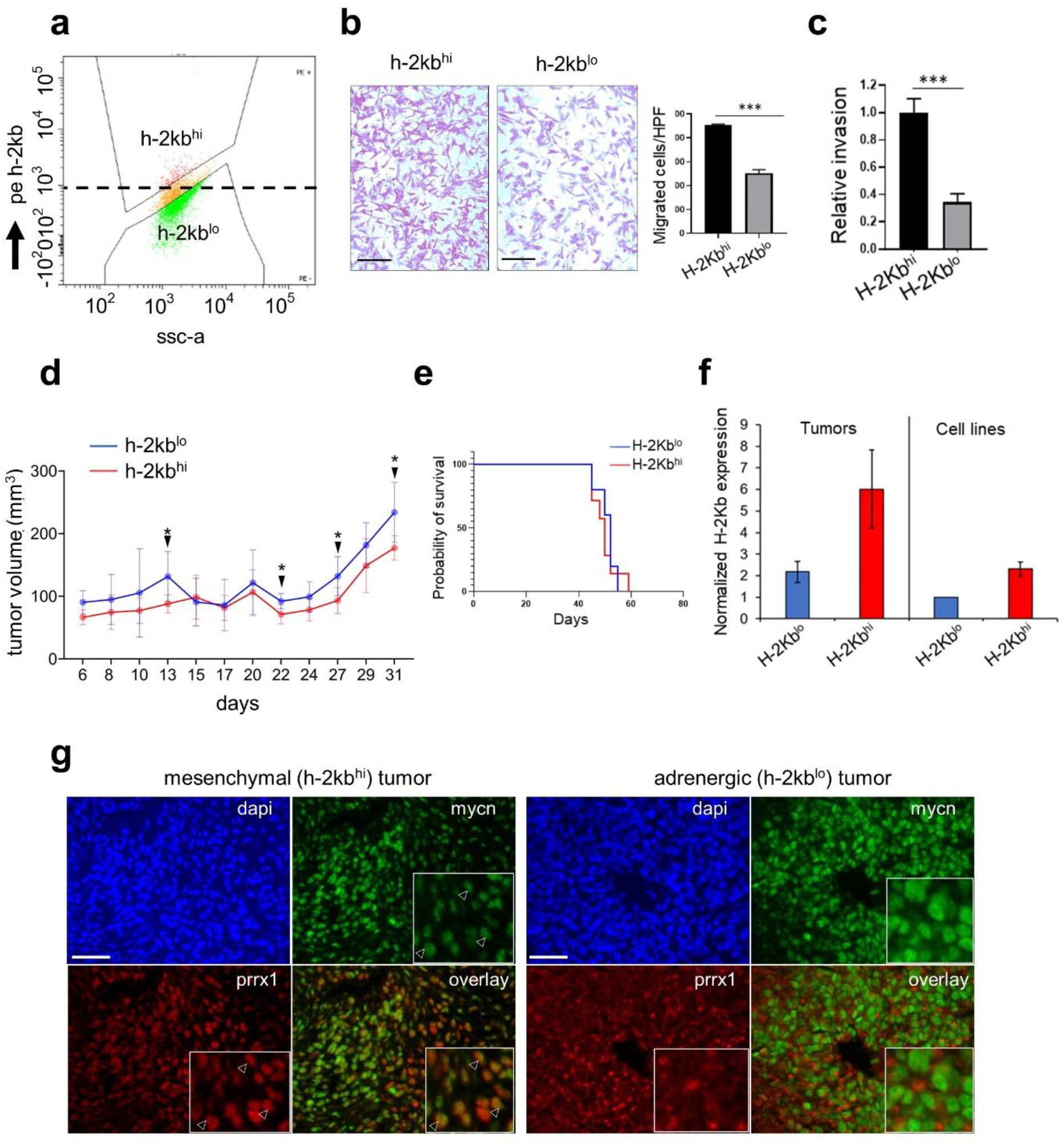
Tumors that arise from NB-9464 H-2Kb^hi^ (mesenchymal) cells show cytotoxic T cell infiltration. **(a)** FACS scatter plot showing gating conditions used for sorting NB-9464 cells into H-2Kb^hi^ and H-2Kb^lo^ populations. X axis represents side scatter (SSC-A); Y axis denotes fluorescence intensity of surface H-2Kb detected using phycoerythrin (PE)-conjugated antibody against H-2Kb. A logscale expression value of 10^3^ was used as the threshold (horizontal dashed line) to gate H-2Kb^hi^ (≥ 10^3^) and H-2Kb^lo^ (< 10^3^) populations. **(b)** *Left*, Bright field images of crystal violet-stained H-2Kb^hi^ and H-2Kb^lo^ cells in transwell migration assays. Scale bars, 100 µm. *Right*, Quantification of migrating cells per high-power field (HPF). Data represent the means ± SD, *n* = 2 biological replicates, \*\*\**P* < 0.001; two-tailed Student’s t-test. **(c)** Quantification of the relative invasiveness of H-2Kb^hi^ and H-2Kb^lo^ cells. Data represent the means ± SD, *n* = 2 biological replicates, \*\*\**P* < 0.001; two-tailed Student’s t-test. **(d)** Tumor volumes in immunocompetent C57BL/6 mice injected subcutaneously with 1 x 10^6^ H-2Kb^lo^ or H-2Kb^hi^ NB-9464 cells. Measurements were started on day 3 after injection and continued three times weekly for up to 50 days or until euthanized due to tumor growth. Graphs represent changes in tumor volume until day 31 (means ± SD; n = 7 per group at all time points) to highlight an earlier onset of tumor formation with H-2Kb^lo^ cells compared to H-2Kb^hi^ cells. Tumors were considered to be established upon reaching a volume of ∼250 mm^3^ (observed between days 31-34 for both H-2Kb^lo^ and H-2Kb^hi^ tumors), following which both tumor types displayed equal increases in tumor growth (data not shown). Tumor onset was defined as the day following which tumor volumes showed a consistent increase (24.3 ± 2.2 days for H-2Kb^lo^ and 27.4 ± 2.1 days for H-2Kb^hi^ cells, *P* < 0.05). Closed arrows refer to the indicated days (13, 22, 27 and 31) on which there were significant differences in tumor volumes (day 13, *P* = 0.02; day 22, *P* = 0.01; day 27, *P* = 0.02 and day 31, *P* = 0.02; *n* = 7 per group at all time points). All *P*-values calculated using the two-tailed Student’s t-test. **(e)** Kaplan-Meier survival analysis of immunocompetent C57BL/6 mice bearing NB tumor xenografts derived from H-2Kb^lo^ (adrenergic) and H-2Kb^hi^ (mesenchymal) cells (50.8 ± 3.7 vs. 49.8 ± 4.8 days; *P* = 0.7; n = 7 per group; log-rank test. **(f)** RT-qPCR analysis of H-2Kb expression in H-2Kb^lo^ and H-2Kb^hi^ NB tumor xenografts and the cell lines used to generate the xenografts. Data represent the means ± SD, *n* = 3 biological replicates. **(g)** Immunofluorescence images of MYCN (green) and Prrx1 (red) expression in representative murine NB xenograft tumors derived from NB-9464 H-2Kb^hi^ (mesenchymal) and H-2Kb^lo^ (adrenergic) cells in immunocompetent syngeneic (C57BL/6) mice. Nuclei are counterstained with DAPI (blue). Insets depict cells with nuclear co-staining of MYCN and Prrx1 (arrowheads) and are exclusively present in the H-2Kb^hi^ mesenchymal tumor. Scale bars, 100 µm, insets 33.3 µm.

**Supplementary Fig. 9.**
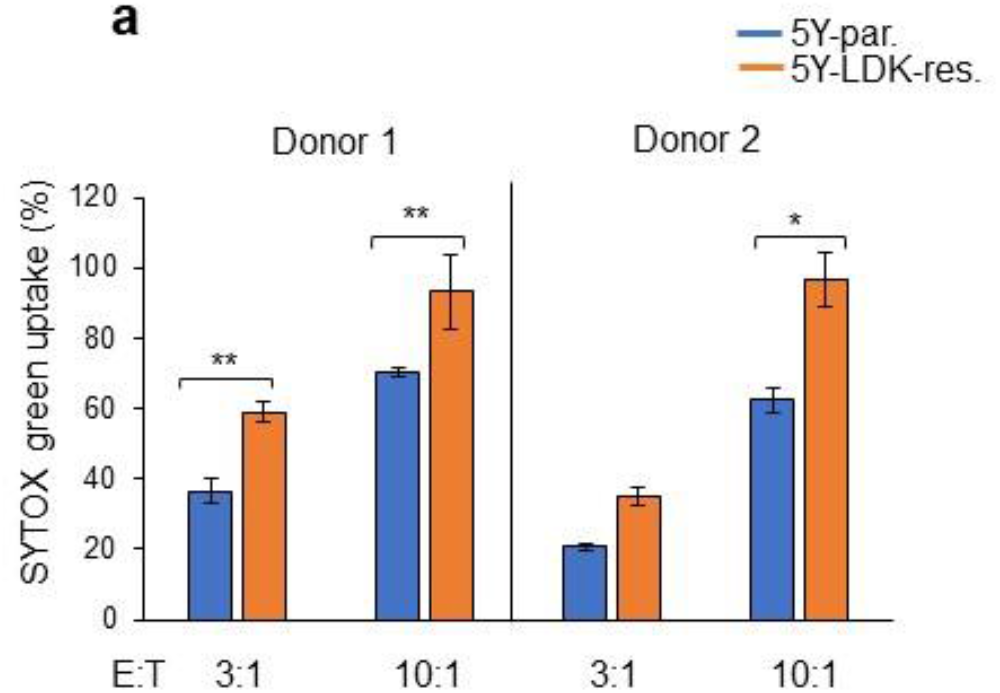
LDK-resistant SH-SY5Y cells are more susceptible to NK-induced cell death. **(a)** Bargraphs showing NK-induced cell death in parental SH-SY5Y and LDK-resistant SH-SY5Y cells ssessed by SYTOX green uptake following 1 hour of co-culture at indicated effector-to-target (E:T) ratios. xperiments were performed in two biological replicates using NK cells harvested from two independent onors. Data represent means ± SD, n = 3 technical replicates. Significance for all results was calculated sing the paired two-tailed Student’s t-test (\**P* < 0.05; **P < 0.01).

## References

1. Yu, A.L. et al. Anti-GD2 antibody with GM-CSF, interleukin-2, and isotretinoin for neuroblastoma. N Engl J Med 363, 1324–34 (2010).

2. Cheung, N.K. & Dyer, M.A. Neuroblastoma: developmental biology, cancer genomics and immunotherapy. Nat Rev Cancer 13, 397–411 (2013).

3. Singh, N. et al. T cells targeting NY-ESO-1 demonstrate efficacy against disseminated neuroblastoma. Oncoimmunology 5, e1040216 (2016).

4. . Park, J.R. et al. Adoptive transfer of chimeric antigen receptor re-directed cytolytic T lymphocyte clones in patients with neuroblastoma. Mol Ther 15, 825–33 (2007).

5. Pule, M.A. et al. Virus-specific T cells engineered to coexpress tumor-specific receptors: persistence and antitumor activity in individuals with neuroblastoma. Nat Med 14, 1264– 70 (2008).

6. Louis, C.U. et al. Antitumor activity and long-term fate of chimeric antigen receptor-positive T cells in patients with neuroblastoma. Blood 118, 6050–6 (2011).

7. Merchant, M.S. et al. Phase I Clinical Trial of Ipilimumab in Pediatric Patients with Advanced Solid Tumors. Clin Cancer Res 22, 1364–70 (2016).

8. Davis, K.L. et al. Nivolumab in children and young adults with relapsed or refractory solid tumours or lymphoma (ADVL1412): a multicentre, open-label, single-arm, phase 1-2 trial. Lancet Oncol 21, 541–550 (2020).

9. Richards, R.M., Sotillo, E. & Majzner, R.G. CAR T Cell Therapy for Neuroblastoma. Front Immunol 9, 2380 (2018).

10. Bernards, R., Dessain, S.K. & Weinberg, R.A. N-myc amplification causes down-modulation of MHC class I antigen expression in neuroblastoma. Cell 47, 667–74 (1986).

11. Burr, M.L. et al. An Evolutionarily Conserved Function of Polycomb Silences the MHC Class I Antigen Presentation Pathway and Enables Immune Evasion in Cancer. Cancer Cell 36, 385–401 e8 (2019).

12. Raffaghello, L. et al. Multiple defects of the antigen-processing machinery components in human neuroblastoma: immunotherapeutic implications. Oncogene 24, 4634–44 (2005).

13. Raffaghello, L. et al. Downregulation and/or release of NKG2D ligands as immune evasion strategy of human neuroblastoma. Neoplasia 6, 558–68 (2004).

14. Castriconi, R. et al. Identification of 4Ig-B7-H3 as a neuroblastoma-associated molecule that exerts a protective role from an NK cell-mediated lysis. Proc Natl Acad Sci U S A 101, 12640–5 (2004).

15. Coughlin, C.M. et al. Immunosurveillance and survivin-specific T-cell immunity in children with high-risk neuroblastoma. J Clin Oncol 24, 5725–34 (2006).

16. Betancur, P.A. et al. A CD47-associated super-enhancer links pro-inflammatory signalling to CD47 upregulation in breast cancer. Nat Commun 8, 14802 (2017).

17. Cohen, M.A. et al. Formation of Human Neuroblastoma in Mouse-Human Neural Crest Chimeras. Cell Stem Cell 26, 579–592 e6 (2020).

18. Asgharzadeh, S. et al. Clinical significance of tumor-associated inflammatory cells in metastatic neuroblastoma. J Clin Oncol 30, 3525–32 (2012).

19. Song, L. et al. Valpha24-invariant NKT cells mediate antitumor activity via killing of tumor-associated macrophages. J Clin Invest 119, 1524–36 (2009).

20. Mao, Y. et al. Targeting Suppressive Myeloid Cells Potentiates Checkpoint Inhibitors to Control Spontaneous Neuroblastoma. Clin Cancer Res 22, 3849–59 (2016).

21. Tran, H.C. et al. TGFbetaR1 Blockade with Galunisertib (LY2157299) Enhances Anti-Neuroblastoma Activity of the Anti-GD2 Antibody Dinutuximab (ch14.18) with Natural Killer Cells. Clin Cancer Res 23, 804–813 (2017).

22. Brodeur, G.M., Seeger, R.C., Schwab, M., Varmus, H.E. & Bishop, J.M. Amplification of N-myc in untreated human neuroblastomas correlates with advanced disease stage. Science 224, 1121–4 (1984).

23. Seeger, R.C. et al. Association of multiple copies of the N-myc oncogene with rapid progression of neuroblastomas. N Engl J Med 313, 1111–6 (1985).

24. Layer, J.P. et al. Amplification of N-Myc is associated with a T-cell-poor microenvironment in metastatic neuroblastoma restraining interferon pathway activity and chemokine expression. Oncoimmunology 6, e1320626 (2017).

25. Wei, J.S. et al. Clinically Relevant Cytotoxic Immune Cell Signatures and Clonal Expansion of T-Cell Receptors in High-Risk MYCN-Not-Amplified Human Neuroblastoma. Clin Cancer Res 24, 5673–5684 (2018).

26. Brandetti, E. et al. MYCN is an immunosuppressive oncogene dampening the expression of ligands for NK-cell-activating receptors in human high-risk neuroblastoma. Oncoimmunology 6, e1316439 (2017).

27. Valentijn, L.J. et al. Functional MYCN signature predicts outcome of neuroblastoma irrespective of MYCN amplification. Proc Natl Acad Sci U S A 109, 19190–5 (2012).

28. Durbin, B.P., Hardin, J.S., Hawkins, D.M. & Rocke, D.M. A variance-stabilizing transformation for gene-expression microarray data. Bioinformatics 18 Suppl 1, S105–10 (2002).

29. Lin, S.M., Du, P., Huber, W. & Kibbe, W.A. Model-based variance-stabilizing transformation for Illumina microarray data. Nucleic Acids Res 36, e11 (2008).

30. McInnes L, H.J. UMAP: Uniform manifold approximation and projection for dimension reduction. arXiv preprint *arXiv:1802.03426*. (2018).

31. . Becht, E. et al. Dimensionality reduction for visualizing single-cell data using UMAP. Nat Biotechnol (2018).

32. Brodeur, G.M. et al. Revisions of the international criteria for neuroblastoma diagnosis, staging, and response to treatment. Journal of Clinical Oncology 11, 1466–1477 (1993).

33. Cohn, S.L. et al. The International Neuroblastoma Risk Group (INRG) classification system: an INRG Task Force report. J Clin Oncol 27, 289–97 (2009).

34. Satija, R., Farrell, J.A., Gennert, D., Schier, A.F. & Regev, A. Spatial reconstruction of single-cell gene expression data. Nat Biotechnol 33, 495–502 (2015).

35. Spranger, S. et al. Up-regulation of PD-L1, IDO, and T(regs) in the melanoma tumor microenvironment is driven by CD8(+) T cells. Sci Transl Med 5, 200ra116 (2013).

36. Wherry, E.J. & Kurachi, M. Molecular and cellular insights into T cell exhaustion. Nat Rev Immunol 15, 486–99 (2015).

37. Boeva, V. et al. Heterogeneity of neuroblastoma cell identity defined by transcriptional circuitries. Nat Genet 49, 1408–1413 (2017).

38. van Groningen, T. et al. Neuroblastoma is composed of two super-enhancer-associated differentiation states. Nat Genet 49, 1261–1266 (2017).

39. Bindea, G. et al. Spatiotemporal dynamics of intratumoral immune cells reveal the immune landscape in human cancer. Immunity 39, 782–95 (2013).

40. Wang, L.L. et al. Augmented expression of MYC and/or MYCN protein defines highly aggressive MYC-driven neuroblastoma: a Children’s Oncology Group study. Br J Cancer 113, 57–63 (2015).

41. Casey, S.C. et al. MYC regulates the antitumor immune response through CD47 and PD-L1. Science 352, 227–31 (2016).

42. Cursons, J. et al. A Gene Signature Predicting Natural Killer Cell Infiltration and Improved Survival in Melanoma Patients. Cancer Immunol Res 7, 1162–1174 (2019).

43. Newman, A.M. et al. Robust enumeration of cell subsets from tissue expression profiles. Nat Methods 12, 453–7 (2015).

44. Hu, X. et al. Landscape of B cell immunity and related immune evasion in human cancers. Nat Genet 51, 560–567 (2019).

45. Ross, R.A., Spengler, B.A. & Biedler, J.L. Coordinate morphological and biochemical interconversion of human neuroblastoma cells. J Natl Cancer Inst 71, 741–7 (1983).

46. Cohn, S.L. et al. Prolonged N-myc protein half-life in a neuroblastoma cell line lacking N-myc amplification. Oncogene 5, 1821–7 (1990).

47. Debruyne, D.N. et al. ALK inhibitor resistance in ALK(F1174L)-driven neuroblastoma is associated with AXL activation and induction of EMT. Oncogene 35, 3681–91 (2016).

48. Chipumuro, E. et al. CDK7 inhibition suppresses super-enhancer-linked oncogenic transcription in MYCN-driven cancer. Cell 159, 1126–1139 (2014).

49. Durbin, A.D. et al. Selective gene dependencies in MYCN-amplified neuroblastoma include the core transcriptional regulatory circuitry. Nat Genet 50, 1240–1246 (2018).

50. Margueron, R. & Reinberg, D. The Polycomb complex PRC2 and its mark in life. Nature 469, 343–9 (2011).

51. Kroesen, M. et al. A transplantable TH-MYCN transgenic tumor model in C57Bl/6 mice for preclinical immunological studies in neuroblastoma. Int J Cancer 134, 1335–45 (2014).

52. Weiss, W.A., Aldape, K., Mohapatra, G., Feuerstein, B.G. & Bishop, J.M. Targeted expression of MYCN causes neuroblastoma in transgenic mice. EMBO J 16, 2985–95 (1997).

53. Clarke, S.R. et al. Characterization of the ovalbumin-specific TCR transgenic line OT-I: MHC elements for positive and negative selection. Immunol Cell Biol 78, 110–7 (2000).

54. Cibrian, D. & Sanchez-Madrid, F. CD69: from activation marker to metabolic gatekeeper. Eur J Immunol 47, 946–953 (2017).

55. Uhrberg, M. The CD107 mobilization assay: viable isolation and immunotherapeutic potential of tumor-cytolytic NK cells. Leukemia 19, 707–9 (2005).

56. Molfetta, R. et al. Regulation of NKG2D Expression and Signaling by Endocytosis. Trends Immunol 37, 790–802 (2016).

57. Qi, W. et al. An allosteric PRC2 inhibitor targeting the H3K27me3 binding pocket of EED. Nat Chem Biol 13, 381–388 (2017).

58. Dongre, A. & Weinberg, R.A. New insights into the mechanisms of epithelial-mesenchymal transition and implications for cancer. Nat Rev Mol Cell Biol 20, 69–84 (2019).

59. Malladi, S. et al. Metastatic Latency and Immune Evasion through Autocrine Inhibition of WNT. Cell 165, 45–60 (2016).

60. Pereira, B.I. et al. Senescent cells evade immune clearance via HLA-E-mediated NK and CD8(+) T cell inhibition. Nat Commun 10, 2387 (2019).

61. Shin, H. & Wherry, E.J. CD8 T cell dysfunction during chronic viral infection. Curr Opin Immunol 19, 408–15 (2007).

62. Kortlever, R.M. et al. Myc Cooperates with Ras by Programming Inflammation and Immune Suppression. Cell 171, 1301–1315 e14 (2017).

63. Sodir, N.M. et al. Endogenous Myc maintains the tumor microenvironment. Genes Dev 25, 907–16 (2011).

64. Bellmeyer, A., Krase, J., Lindgren, J. & LaBonne, C. The protooncogene c-myc is an essential regulator of neural crest formation in xenopus. Dev Cell 4, 827–39 (2003).

65. Rada-Iglesias, A. et al. Epigenomic annotation of enhancers predicts transcriptional regulators of human neural crest. Cell Stem Cell 11, 633–48 (2012).

66. Zeid, R. et al. Enhancer invasion shapes MYCN-dependent transcriptional amplification in neuroblastoma. Nat Genet 50, 515–523 (2018).

67. Chang, C.H., Hammer, J., Loh, J.E., Fodor, W.L. & Flavell, R.A. The activation of major histocompatibility complex class I genes by interferon regulatory factor-1 (IRF-1). Immunogenetics 35, 378–84 (1992).

68. Unterholzner, L. et al. IFI16 is an innate immune sensor for intracellular DNA. Nat Immunol 11, 997–1004 (2010).

69. Spel, L. et al. Nedd4-Binding Protein 1 and TNFAIP3-Interacting Protein 1 Control MHC-1 Display in Neuroblastoma. Cancer Res 78, 6621–6631 (2018).

70. Sen, T. et al. Targeting DNA Damage Response Promotes Antitumor Immunity through STING-Mediated T-cell Activation in Small Cell Lung Cancer. Cancer Discov 9, 646–661 (2019).

71. Ennishi, D. et al. Molecular and Genetic Characterization of MHC Deficiency Identifies EZH2 as Therapeutic Target for Enhancing Immune Recognition. Cancer Discov 9, 546–563 (2019).

72. Zingg, D. et al. The Histone Methyltransferase Ezh2 Controls Mechanisms of Adaptive Resistance to Tumor Immunotherapy. Cell Rep 20, 854–867 (2017).

73. Peng, D. et al. Epigenetic silencing of TH1-type chemokines shapes tumour immunity and immunotherapy. Nature 527, 249–53 (2015).

74. Xiao, L. et al. Adoptive Transfer of NKG2D CAR mRNA-Engineered Natural Killer Cells in Colorectal Cancer Patients. Mol Ther 27, 1114–1125 (2019).

75. Schindelin, J. et al. Fiji: an open-source platform for biological-image analysis. Nat Methods 9, 676–82 (2012).

76. Kamentsky, L. et al. Improved structure, function and compatibility for CellProfiler: modular high-throughput image analysis software. Bioinformatics 27, 1179–80 (2011).

77. Love, M.I., Huber, W. & Anders, S. Moderated estimation of fold change and dispersion for RNA-seq data with DESeq2. Genome Biol 15, 550 (2014).

78. Zhu, X. et al. Single-Cell Clustering Based on Shared Nearest Neighbor and Graph Partitioning. Interdiscip Sci 12, 117–130 (2020).

79. Russo, P.S.T. et al. CEMiTool: a Bioconductor package for performing comprehensive modular co-expression analyses. BMC Bioinformatics 19, 56 (2018).

80. Yu, G., Wang, L.G., Han, Y. & He, Q.Y. clusterProfiler: an R package for comparing biological themes among gene clusters. OMICS 16, 284–7 (2012).

81. Alexopoulos, E.C. Introduction to multivariate regression analysis. Hippokratia 14, 23–8 (2010).

82. Das, S. & Bansal, M. Variation of gene expression in plants is influenced by gene architecture and structural properties of promoters. PLoS One 14, e0212678 (2019).

83. Debruyne, D.N. et al. BORIS promotes chromatin regulatory interactions in treatment-resistant cancer cells. Nature 572, 676–680 (2019).

